# Loss of hepatic Lgr4 and Lgr5 promotes nonalcoholic fatty liver disease

**DOI:** 10.1101/2021.11.22.469602

**Authors:** Enrica Saponara, Carlos Penno, Meztli L. Matadamas Guzmán, Virginie Brun, Benoit Fischer, Margaret Brousseau, Zhong-Yi Wang, Peter ODonnell, Jonathan Turner, Alexandra Graff Meyer, Laura Bollepalli, Giovanni d’Ario, Guglielmo Roma, Walter Carbone, Vanessa Orsini, Stefano Annunziato, Michael Obrecht, Nicolau Beckmann, Chandra Saravanan, Arnaud Osmont, Philipp Tropberger, Shola Richards, Christel Genoud, Alexandra Aebi, Svenja Ley, Iwona Ksiazek, Florian Nigsch, Luigi Terraciano, Tewis Bouwmeester, Jan Tchorz, Heinz Ruffner

## Abstract

**Background & Aims:** The Rspo-Lgr4/5-Znrf3/Rnf43 module is a master regulator of hepatic Wnt/β-catenin signaling and metabolic zonation, but its impact on nonalcoholic fatty liver disease (NAFLD) remains unclear. We studied whether liver-specific loss of the Wnt/β-catenin modulators Leucine-Rich Repeat-Containing G Protein-Coupled Receptor 4/5 (Lgr4/5) promotes nonalcoholic fatty liver disease (NAFLD).

**Methods:** Mice with liver-specific deletion of both receptors Lgr4/5 (Lgr4/5dLKO) were fed with normal diet (ND) or high fat diet (HFD). Livers of these mice were analyzed for lipid and fibrotic content by tissue staining and immunohistochemistry (IHC), and lipoproteins, inflammation and liver enzyme markers were measured in blood. Mechanistic insights into hepatic lipid accumulation were obtained by using *ex vivo* primary hepatocyte cultures derived from the Lgr4/5dLKO mice. Lipid analysis of mouse livers was performed by mass spectrometry (MS)-based untargeted lipidomic analysis.

**Results:** We demonstrated that liver-specific ablation of Lgr4/5-mediated Wnt signaling resulted in hepatic steatosis, impaired bile acid (BA) secretion and predisposition to liver fibrosis. Under HFD conditions, we observed progressive intrahepatic fat accumulation, developing into macro-vesicular steatosis. Serum lipoprotein levels in HFD-fed Lgr4/5dLKO mice were decreased, rather than increased, suggesting that accumulation of fat in the liver was due to impaired lipid secretion by hepatocytes. Our lipidome analysis revealed a severe alteration of several lipid species in livers of Lgr4/5dLKO mice, including triacylglycerol estolides (TG-EST), a storage form of bioactive free fatty acid (FA) esters of hydroxy FAs (FAHFAs).

**Conclusions:** Loss of hepatic Wnt/β-catenin activity by Lgr4/5 deletion led to deregulation of lipoprotein pathways, loss of BA secretion, intrinsic alterations of lipid homeostasis and the onset of NAFLD.

**Lay summary:** The Wnt/β-catenin pathway plays an important role during development and tissue homeostasis. Loss of Wnt/β-catenin activity in mouse liver leads to loss of liver zonation, but the impact on nonalcoholic fatty liver disease (NAFLD) remains unclear. We show that livers of mice developed steatosis upon deletion of the positive pathway regulators Lgr4/5. Livers of knock-out (KO) mice exhibited altered lipid composition due to impaired lipid secretion. Furthermore, livers of these mice developed a nonalcoholic steatohepatitis (NASH)-like phenotype and fibrotic features derived from activated hepatic stellate cells. Our data demonstrate a protective role of Wnt/β-catenin pathway activity towards the development of NAFLD.

**Highlights:**

- Abrogation of hepatic Wnt/β-catenin activity and liver zonation upon Lgr4/5 deletion in mice led to hepatic steatosis.
- Liver fat accumulation was caused by impaired lipid secretion from hepatocytes.
- Steatotic livers contained increased levels of diverse lipid species, including polyunsaturated fatty acids and triglycerol-estolides.
- These data confirmed that a decrease in Wnt/β-catenin signaling led to the development of nonalcoholic fatty liver disease (NAFLD) in mice.

## Introduction

The liver consists of repetitive hexagonal structures called lobules. Hepatocytes in each lobule are divided into three zones, containing periportal (zone 1), midzonal (zone 2) and pericentral (zone 3) hepatocytes. Liver function critically depends on complementary metabolic tasks distributed to hepatocytes alongside the porto-central blood flow. Examples for the spatial segregation of opposing metabolic tasks are the oxidative breakdown of FAs (zone 1) and lipid biosynthesis (zone 3), or gluconeogenesis (zone 1) and glycolysis (zone 3). Disruption of metabolic liver zonation is therefore implicated in several metabolic disorders, such as diabetes and nonalcoholic fatty liver disease (NAFLD) [1, 2].

NAFLD comprises a spectrum of liver conditions including simple steatosis, nonalcoholic steatohepatitis (NASH), fibrosis and cirrhosis, progressively leading to end-stage liver disease or hepatocellular carcinoma. Despite great progress in understanding the complexity of NAFLD, the understanding of the etiology and the pathophysiological mechanisms implicated in liver damage remains incomplete. One of the earliest hallmarks of NAFLD is the intrahepatic accumulation of neutral lipids concomitant with the alteration of lipid concentration in serum, a phenomenon known as dyslipidemia [3]. This imbalance is often attributable to altered hepatic lipid or BA synthesis, import or export, or alterations in intracellular cholesterol biosynthesis [4, 5]. However, the instructive cues for these processes are not fully understood.

A centro-portal Wnt/β-catenin activity gradient has been demonstrated to establish and maintain metabolic liver zonation. Moreover, increasing evidence suggests that Wnt/β-catenin signaling plays a role in both exogenous and endogenous lipid trafficking, regulates the expression BA biosynthesis enzymes in hepatocytes [6, 7], and is required for proper bile canaliculi morphogenesis [8, 9]. When Wnt ligands bind their cognate Frizzled receptor and the Lrp5/6 co-receptor, phosphorylation of Lrp5/6 stabilizes β-catenin by decreasing activity of the β-catenin destruction complex. Stabilized β-catenin accumulates in the cytoplasm and can now enter the nucleus, where it binds to TCF family transcription factors and enhances the transcription of β-catenin target genes [10], including many pericentral metabolic enzymes, such as Cyp2e1 and Cyp2a1 or the tight junction (TJ) protein Claudin-2 (CLDN2) [11]. Consequently, loss of Wnt/β-catenin signaling abrogates the expression of pericentral metabolic genes while increasing periportal metabolic genes, whereas pathway activation caused the opposite phenotype [12–14]. Roof plate-specific spondin (Rspo)1–4 ligands potentiate Wnt/β-catenin signaling following binding to their receptors Lgr4–6 by clearing the cell-surface transmembrane E3 ubiquitin ligases Zinc and ring finger 3 (Znrf3) and its homologue Ring finger protein 43 (Rnf43), which promote Wnt receptor turnover at the plasma membrane [15–18]. Complete loss of hepatic Wnt/β-catenin signaling in mice with liver-specific deletion of Lgr4/5 showed that the Rspo-Lgr4/5-Znrf3/Rnf43 module is a critical regulator, rather than just a potentiator, of hepatic Wnt/β-catenin activity and metabolic zonation [19]. Moreover, combined deletion of *Znrf3/Rnf43* or Rspo injections expanded the pericentral metabolic program throughout the liver, reprogramming periportal into pericentral hepatocytes and thus disrupting metabolic liver zonation. However, the implication of the Rspo-Lgr4/5-Znrf3/Rnf43 module on NAFLD is unclear.

A recent report suggests that hepatocyte-specific loss of Znrf3/Rnf43 leads to a NASH-like phenotype in ND-fed mice because of altered lipid metabolism. The authors propose that the mutant mice exhibit altered hepatocyte regeneration that predisposes to hepatocellular carcinoma [20]. In addition, hepatocyte-specific expression of an activated β-catenin transgene resulted into pronounced pericentral steatosis under HFD conditions, showing diet-induced obesity, systemic insulin resistance and increased hepatic expression of glycolytic and lipogenic genes, whereas animals fed with ND had normal liver histology and body weight [21]. In contrast, we have recently shown that combined deletion of Znrf3/Rnf43 and the resulting increase in Wnt/β-catenin signaling did not lead to the spontaneous development of NAFLD but promoted uncontrolled hepatocyte proliferation and tumor formation within an observation period of one year [22]. Our findings are in line with previous work suggesting that liver-specific loss of Wnt/β-catenin signaling, rather than its increase leads to NAFLD and NASH [21, 23, 24].

To clarify the controversial role of the Rspo-Lgr4/5-Znrf3/Rnf43 module in NAFLD, we studied lipid metabolism and NAFLD formation in mice with liver-specific deletion of Lgr4/5. Mice of different age were analyzed for steatotic and fibrotic manifestation under ND and HFD conditions, and *ex vivo* hepatocyte cultures from the KO and WT mice were utilized to study lipid uptake and storage and BA secretion. Our results demonstrated that loss of Wnt/β-catenin signaling upon deletion of Lgr4/5 in mouse liver led to a NASH-like phenotype characterized by steatosis, increased inflammation and fibrosis upon HFD. Several lipid species were increased in livers of KO mice, and serum lipids were decreased.

## Materials and Methods

### Mouse model generation and animal experimentation, *ex vivo* assays

All animal experimentation was conducted in Basel, Switzerland, in accordance with the Swiss Animal Welfare Legislation and authorized by the cantonal veterinary office of Basel city.

Liver-specific deletion of both Lgr4 and Lgr5 receptors was achieved as previously described [19]. Briefly, mouse models for conditional KO of both Lgr4 (Lgr4lox) and Lgr5 (Lgr5lox) were crossed with *Alb-Cre* mice (Lgr4/5dLKO). *Alb-Cre* negative, *Lgr4/5^fl/fl^* mice were utilized as controls. Unless differently specified, both male and female animals were used in this study.

#### HFD

For 3 months- and 6 months-old time points, mice were housed with *ad libitum* access to water and food until the age of 2 and 5 months, respectively. Afterwards they were fed with HFD, 45 % kcal (Kliba/NAFAG, #2126) for 4 consecutive weeks (1 month). Age-matched animals were maintained on normal diet (ND).

#### LDL-Phrodo assay

Primary hepatocytes were kept overnight in basal medium. On the following day, LDL-Phrodo red (Life Technologies, #I34360) was added to each well at the final concentration of 10 μg/ml. Imaging and optical density were acquired after 5, 10, 15, 20 and 24 hours with Arrayscan XTI (Cellomics, Thermo Fisher Scientific). After the final measurement, cells were washed with PBS and incubated 1 minute with BODIPY FL working solution at 20 μg/ml together with the nuclear dye Hoechst.

#### CLF assay

After seeding, cells were incubated in attachment medium for 4 hours. Afterward, ice-cold matrigel was added at a final concentration of 250 μg/ml. Cells were cultured over a period of 7 days in basal medium. Matrigel was renewed every second day. At day 7, cells were incubated for 25 min at 37°C with Cholyl-L-Lysyl-Fluorescein (CLF, Corning #33288) at a final concentration of 5 mM. CLF-positive canaliculi were imaged utilizing Arrayscan XTI (Cellomics, Thermo Fisher Scientific).

### Lipid droplet analysis

After isolation, cells were cultured in basal media over 24 hours. Cells were fixed with 10% PFA and afterward stained with BODIPY FL. Lipid droplet-size quantification was performed with the cell imaging software Imaris (Oxford instrument, UK).

#### RNA isolation and library preparation

Total RNA was isolated from mouse livers with Qiazol (Invitrogen, #79306) and purified with the miRNeasy kit (Qiagen, #217004). The amount of RNA was quantified with the Agilent RNA 6000 Nano Kit (Agilent Technologies, #5067-1511). RNA-sequencing libraries were prepared from mouse liver total RNA using the Illumina TruSeq Stranded Total RNA sample preparation protocol following the manufacturer’s instructions.

#### RNA sequencing

RNA sequencing and related data analysis were performed as described in [19]. Differential expression among Lgr4/5dLKO and control mice was visualized as volcano plots with the TIBCO Spotfire software. RNA-Seq raw data are available at the NIH Sequence Read Archive with the following ID: PRJNA670351.

#### RNA extraction, reverse transcription and quantitative PCR (qPCR)

Total liver RNA was isolated using the RNeasy mini kit including on-column DNase digestion according to the manufacturer’s instructions (Qiagen). Two micrograms of RNA from each tissue sample were reverse-transcribed using the high-capacity cDNA reverse transcription kit (Applied Biosystems). The resulting cDNA products were diluted 1:20 and subjected to qPCR reactions using TaqMan reagents. qPCR reactions were conducted on an ABI Prism 7900HT Sequence Detection System (Applied Biosystems). The threshold crossing value (Ct) was determined for each transcript and normalized to the internal control transcript Gapdh. The relative quantification of each mRNA species was assessed using the comparative Ct method. Statistical analysis was performed using the Student’s t-test with the Graphpad Prism software (Graphpad Software 8.1.2). Primers utilized in this study were Cdh1, Mm01247357_m1; Ctnnb1, Mm00483039_m1; Axin2, Mm00443610_m1.

#### Pathway enrichment analysis for differentially expressed genes

The clusterProfiler [25] and ReactomePA [26] R packages were used to perform Reactome pathway enrichment analysis for upregulated and downregulated genes in Lgr4/5dLKO mice. Fisher’s exact test followed by the Benjamini-Hochberg correction was performed for statistical analysis, and an adjusted *P* < 0.05 was set as the cutoff criterion. A gene was declared differentially expressed if a difference observed in expression between two experimental conditions was statistically significant (adjusted p-value (*P*) < 0.05). Log_2_ fold change of expression was used to define up- or down-regulation.

#### Statistics

For analysis of serum biochemistry (ALT, AST, t-BA), HTRF assays (IL6, TNFα), CLF assay, BODIPY quantification, CDH1, AXIN2, and CTNNB1, gene expression and elastography between two groups, a twotailed t test was performed. For analysis of LDL-Phrodo uptake over time and serum levels of t-Chol, c-HDL and TG, a two-way ANOVA was performed. A P < 0.05 was considered significant, and plots display mean ± standard deviation (SD) in case of biological replicates, ± standard error of the mean (SEM) for technical replicates. All statistical analyses and graph generation were performed using the GraphPad Prism 8.1.2 software. For RNA sequencing analysis, the method utilized is described in [19]. Briefly, Limma R package from the Bioconductor suite was used to identify genes differentially expressed between the conditions of interest. P values were adjusted for multiple hypothesis using the Benjamini-Hochberg correction. Genes were defined differentially expressed if P ≤ 0.01 with at least a two-fold change in either direction.

#### Blood sample preparation

After each experiment, animals were euthanized with CO2, and blood was collected from the vena cava. Serum was obtained by centrifuging blood at 10’000 g for 90 sec into a Microtainer® Tube, 400 μL to 600 μL, SST™ Gel Serum Separator Additive, Clear (BD #365967), and kept at −20°C until further processing.

#### Fast protein liquid chromatography (FPLC)-based size-exclusion

50-100 μL of serum per samples were pooled and utilized for FPLC measurement. FPLC was conducted using a BioLogic DuoFlow FPLC System - Two column (BioRad), Superose 6 Increase, 10/300GL, FPLC column (GE Healthcare, #29-0915-96), BioLogic BioFrac Fraction Collector (BioRad), blood bank saline. To measure total cholesterol and triglycerol, these reagents were used: SpectraMax M2 Spectrophotometer (Molecular Devices), Total Cholesterol Kit (Wako Diagnostics, Cholesterol E), L-Type Triglycerol M (Wako Diagnostics, Enzyme Color R1 & R2), 96-well, half-area plate (Corning #3690).

#### Primary hepatocyte purification, long-term culture

Perfusion procedures were performed at 37°C, 7 ml/min constant flow rate with the Harvard mini-peristaltic pump. Livers were initially cleared from blood utilizing 35-50 ml/mouse of pre-perfusion solution consisting of 1X HBSS -/-, 0.5 mM EGTA, 10 mM HEPES. After 5 min, the pre-perfusion solution was substituted with perfusion solution: DMEM-F12, 3 mM CaCl2, PenStrep, 20 mM HEPES, Collagenase (120 CDU/ml, Roche #11088793001). The liver was perfused for the next 10 min, removed from the abdominal cavity and placed into a 10-cm dish with ice-cold wash solution (DMEM/F12 supplemented with PenStrep and 10% FBS). Liver lobes were teared apart with forceps and hepatic cells released into wash solution.

Cells were collected and 2x washed in wash solution by centrifugation at 50 g for 5 min, 4°C, then suspended in attachment media consisting of Williams E medium without phenol red, PenStrep, 10% FBS, 1:10000 insulin (1.7 mM), Glutamax, 0.3 mM dexamethasone, 10 mM HEPES. Cells were further purified on a 21.6 ml Percoll (GE Healthcare, #17-0891-02) gradient supplemented with 2.4 ml of 10x HBSS and finally washed 2x in 25 ml attachment medium by centrifugation at 60 g for 3 min, 4°C. 220’000 cells/well were distributed on 24-well Collagen 1-coated plates (BD, #356408) and incubated at 37°C, 5% CO2. The basal media utilized in all conditions consisted of Williams E medium (Sigma #W1878), PenStrep, 1:10000 insulin (1.7 mM), Glutamax, 1:10000 Dexamethasone (0.3 mM), 10 mM HEPES.

#### Glucose and insulin tolerance tests (GTT, ITT)

Mice were fasted for 17 and 6 hours, respectively, and baseline blood glucose levels were measured in tail-vein blood using an Accu-Chek Compact plus glucometer (Roche Diagnostics). Glucose (3 g/kg body weight) or insulin (0.75 U/kg body weight) were injected intraperitoneally (i.p). Blood glucose levels were measured prior to and at 15, 30, 45, 60, 90 and 120 min after glucose or insulin injection as previously described [27].

#### Ultrasound elastography

Acquisition: Animals were anesthetized using 2.0% isoflurane (Abbott, Cham, Switzerland) in 100% O2 and placed on an electrical warming pad to maintain body temperature at 37°C throughout the assessment. The anesthesia was maintained during the image acquisition by delivering the anesthetic via a nose cone, and the eyes were protected with Viscotears® Augen-Gel creme. The animals were positioned in a supine position. Abdomen was shaved and mouse fixed with tape on upper and lower part of the body. All examinations were performed according to a standardized protocol with an ultrasound (US) system (Aixplorer; SuperSonic Imagine, Aix-En-Provence, France) equipped with a 25 MHz (SL25-15) superficial linear transducer. The transducer was fixed to a micro-manipulator and positioned precisely in the transversal plane. The examination began with gray-scale B-mode to visualize sternum and liver underneath. Afterwards, three measurements in the shear wave elastography (SWE) mode were made in three different places from sternum to stomach. US gel (Aquasonic 100, Parker Laboratories Inc., Fairfield NJ, U.S.A.) was placed between the skin and the US transducer. During the SWE assessment, care was taken not to put any pressure to the liver by the transducer. The liver elasticity (kPa) was measured in SWE mode by placing the region of interest (ROI) at this thickest part of the liver in every slice (Total 3 value).

#### Immunohistochemistry

Three μm formalin-fixed, paraffin-embedded (FFPE) mouse liver sections were stained using standard protocols, stained by hematoxylin and eosin (H&E) and picro-sirius red, and immunofluorescence was conducted to detect CDH1 by using the mouse anti-Cadherin 1 antibody Clone 36 (BD, #610181), anti-IBA1 (Abcam, #ab178846), Rat anti-KRT19 (Developmental Studies Hybridoma Bank at #TROMA-III; RRID: AB_2133570), Rabbit anti-alpha Smooth Muscle Actin (αSMA) antibody [EPR5368] (Abcam, #ab124964). Seven μm thick-OCT-embedded tissue or primary hepatocytes ware stained by BODIPY FL (Life technologies, #D2184), rabbit anti-ZO-1 (Invitrogen, #617300). Digital images were acquired and analyzed with the Aperio ScanScope XT system (Leica Biosystems, Germany) or with a Zeiss LSM 700 (Zeiss, Germany) confocal microscope using the Zen 2011 SP3 (black edition) software. Lipid droplet size quantification was performed with the cell imaging software Imaris (Oxford instrument, UK).

#### IL6 and TNFα HTRF assays

IL6 and TNFα were measured in sera of mice utilizing the mouse IL6 assay (#62MIL06PEG) and mouse TNFα assay (#62CTNFAPEG) according to Cisbio Bioassays vendor protocols.

#### Lipoprotein and BA determination

AST, ALT, t-Chol, c-HDL and TG were measured in sera of mice utilizing the SPOTCHEM II KENSHIN-2 kit (AxonLab #10013990). t-BA were measured utilizing the fluorimetric bile acid assay kit (Sigma, #MAK309) according to the manufacturer’s protocol.

#### Ultra performance liquid chromatography (UPLC)-Q exactive orbitrap/mass spectrometry (MS)-based untargeted lipidomics analysis

Lipid extraction was done in 5 mg of liver resections. A volume of 800 μL of chloroform:methanol (2:1 v/v) containing 15 μL internal standards (ISTDs, EquiSPLASH, Avanti Polar Lipids) were homogenized using the bead beating device (Precellys, Rockville, MD) maintained at 4°C. After homogenization, samples were mixed for 30 min at 4°C and centrifuged at 17500 RCF for 10 min at 4°C. Supernatants were collected and 100 μL of water was added to induce phase separation. Samples were gently mixed for 30 min at 4°C. The lower phase was collected and evaporated under nitrogen. Samples were reconstituted in 100 μL of isopropanol. A pooled sample (QC sample) was generated to assess analytical quality, and an extraction blank sample was generated to determine contaminant masses derived from reagents and materials.

The LC system consisted of a Dionex Ultimate 3000 RS coupled with a HESI probe to a Q Exactive Orbitrap mass spectrometer. Lipids were separated on an Accucore UPLC C30 column (150 × 2.1 mm; 2.6 μm) (Thermo Fisher Scientific). The column was maintained at 50°C at a flowrate of 0.3 mL/min. The mobile phases consisted of (A) 60:40 (v/v) water:acetonitrile with ammonium formate (5 mM) and formic acid (0.1%) and (B) 80:20 (v/v) isopropanol:acetonitrile with ammonium formate (5 mM) and 0.1% formic acid. A sample volume of 5 μL was used for the injection. The separation was conducted under the following gradient: 0 min 25% (B); 20 min 86% (B); 22 min 90 % (B); 24 min 99% (B); 26 min 99 % (B); 32 min 25% (B). Sample temperature was maintained at 4°C. MS1 and MS/MS (data-dependent MS/MS) data in positive and negative ion modes were independently acquired. For the full MS, the automatic gain control (AGC) target was set as 1 × 10^6^ and the maximum injection time was set as 250 ms. In the data-dependent MS/MS the AGC target was set as 1 × 10^5^ and the maximum injection time was set as 120 ms. The source parameters were as follows: 1) in ESI(+):sheath gas pressure, 60; aux gas flow, 3; spray voltage, 4.0 kV; capillary temperature, 400°C; 2) ESI(-): sheath gas pressure, 40; aux gas flow, 2; spray voltage, 3.5 kV; capillary temperature, 450°C; aux gas heater temperature, 390°C. MS1 mass range, m/z 200–1500; MS1 resolving power, 35,000 FWHM (m/z 200); number of data dependent scans per cycle: 10; MS/MS resolving power, 17,500 FWHM (m/z 200). The data-dependent MS/MS mode top N was 10; the normalized collision energy values were 15, 25 and 35 for both positive and negative; and the isolation window was m/z 1.0. The instrument was tuned using positive and negative ion mode calibration solutions (Thermo Fischer Scientific). The shorthand notation system and nomenclature of lipids used are the same as published in [28].

#### Lipid data analysis and lipid identification

The “raw” format files were also converted to “ABF” format using an ABF converter. MS-DIAL 4.48 (lipidomics mode) was used for peaks exaction, peaks alignment, deconvolution analysis, and identification [28]. Parameter settings were as follows. Data collection: MS1/MS2 tolerance, 0.01/0.05 Da; minimum peak height, 5×10^6^ and 1×10^6^ of amplitude for positive and negative ion mode, respectively. Identification: accurate mass tolerance MS1/MS2, 0.01/0.05 Da., score cut off, 90%. Alignment: retention time tolerance, 0.1 min, MS1 tolerance, 0.01 Da. All of the lipid annotations produced with negative and positive acquisition were merged with in-house built scripts in R (version 3.6.3). Further data processing including missing values imputation, logarithmic transformation (base 2), normalization (based on EquiSPLASH and scaling methods), and group comparisons (empirical Bayes moderated t-tests) were performed in R (version 3.5.3) with packages Metabolomics and NormalizeMets [29]. The detailed nomenclature for the lipid classes identified can be found at http://prime.psc.riken.jp/compms/msdial/lipidnomenclature.html.

#### Transmission Electron Microscopy (TEM)

The mouse liver was cut into 2-3 mm^3^ cubes and immediately immersed in a fixation solution containing 2.5% glutaraldehyde (Electron Microscopy Science) in 0.1 M cacodylate buffer pH 7.4 for 1 hour at room temperature and then overnight at 4°C. After five washes in 0.1 M cacodylate buffer (pH 7.4), a post-fixation in 1% osmium tetroxide in 0.1 M cacodylate buffer (pH7.4) was done. After five washes in ddH2O and dehydration steps in graded alcohol series, the mouse liver pieces were embedded in Embed 812 epoxy resin hard (Electron Microscopy Science) for 12 h and polymerized at 60°C during 48 h. For TEM analysis, a region of interest was selected under bright field. After trimming, silver/gray thin sections (50 nm thickness) were collected on formvar-coated single-slot copper grids (Electron Microscopy Science). After post-staining with 1% uranyl acetate and lead citrate (6 min, each), images were recorded using a FEI Tecnai Spirit (FEI Company) operated at 120 keV using a side-mounted 2K × 2K CCD camera (Veleta, Olympus).

## Results

### Lgr4/5dLKO mice displayed progressive hepatic lipid accumulation and altered serum lipoprotein levels

We previously showed that mice with an Albumin-Cre-mediated (*Alb-Cre; Lgr4/5^fl/fl^*) combined deletion of Lgr4 and Lgr5 in hepatocytes and biliary cells (Lgr4/5dLKO) exhibited compromised metabolic zonation and liver regeneration potential [19]. The current work focused on the consequences of loss of Wnt-dependent metabolic zonation in mouse livers. Specifically, we investigated the liver and serum phenotypes in Lgr4/5dLKO and WT mice at different age. Livers of 3 months-old ND-fed Lgr4/5dLKO mice appeared histologically intact (Fig. 1A). Serum total cholesterol (t-Chol) was reduced in Lgr4/5dLKO compared to control (*Alb-Cre* negative, *Lgr4/5 ^fl/fl^*) mice, whereas cholesterol-high density lipoprotein (c-HDL) and triglycerol (TG) levels were similar (Fig. 1B). t-Chol reduction was modest but significant, since mouse serum t-Chol levels measured over a period of 20 weeks are rather stable [30]. TG levels were similar in the two strains, suggestive of a latent dyslipidemia in Lgr4/5dLKO animals. No abnormality concerning glucose (GTT) and insulin tolerance test (ITT) was detected in ND-fed Lgr4/5dLKO mice (Fig. S1A). To elucidate how Lgr4/5dLKO mice reacted to an obesogenic diet, we exposed 2 months-old Lgr4/5dLKO and control mice to a moderate obesogenic 45 % kcal HFD for 4 weeks (3 months endpoint) compared to mice fed with ND. All Lgr4/5dLKO mice developed mild and scattered liver steatotic foci upon HFD, in contrast to control animals (Fig. 1A). Lgr4/5dLKO mice on HFD showed a marked reduction in serum t-Chol and c-HDL levels compared to control animals on HFD, whereas TG levels were similar (Fig. 1B). Of note, the TG/c-HDL ratio is a diagnostic surrogate of NAFLD [31]. Increased aspartate aminotransferase (AST) levels in serum of 3 months-old HFD-fed Lgr4/5dLKO indicated a potential mild degree of comprised hepatocyte function compared to control animals, whereas alanine aminotransferase (ALT) and total bile acids (t-BA) remained unaltered (Fig. 1C). Cholesterol and TGs are transported in lipoproteins in the blood. To investigate whether lipoprotein secretion was altered in our Lgr4/5dLKO mice, we performed FPLC-based size-exclusion chromatography on murine sera. In pooled sera of 3 months-old ND-fed female and male Lgr4/5dLKO mice, the fraction comprising very low density lipoprotein (VLDL)-cholesterol was increased, while the HDL-cholesterol fraction was modestly reduced (Fig. 1D). Interestingly, analogous to NAFLD patients [32], ND-fed Lgr4/5dLKO mice also displayed a remarkable increase in serum VLDL-triglycerol (TG) content compared to control animals (Fig. 1D).

**Figure 1.**
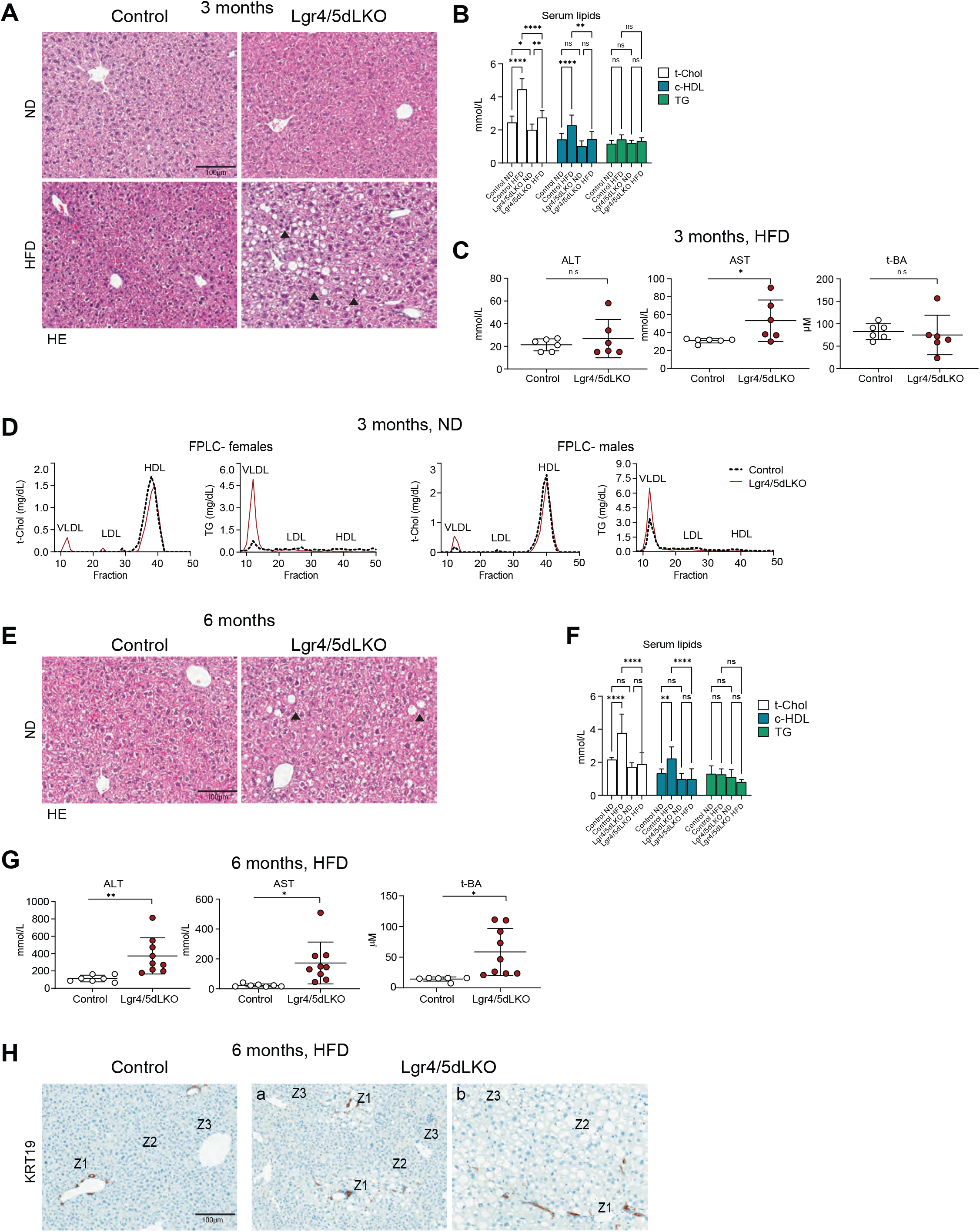
Steatosis, serum lipoprotein imbalance and serum liver enzymes in Lgr4/5dLKO mice. **A.** HE staining on liver sections of 3 months-old animals subjected to ND or 4 weeks HFD, respectively. HFD-fed Lgr4/5dLKO animals developed focal steatosis (arrowheads). **B.** Serum t-Chol levels in ND-fed 3 months-old Lgr4/5dLKO were lower compared to WT control animals, whereas c-HDL and TG levels were similar; in 3 months-old HFD-fed Lgr4/5dLKO mice, t-Chol and c-HDL levels were lower compared to WT animals, whereas TG levels remained unchanged (ND control n=18, ND Lgr4/5dLKO n=8, HFD control n=6, HFD Lgr4/5dLKO n=6; two way ANOVA, **** <0.0001, ** < 0.005, * <0.05). **C**. ALT, AST and t-BA measured in sera of 3 months-old animals subjected HFD (n=6; t test, *= P < 0.05). Note the elevation of AST levels may indicate signs of compromised hepatocyte function. **D.** FPLC analysis on pooled sera of 3 months-old ND-fed mice (female controls n=2, female Lgr4/5dLKO n=3, male controls n=3, male Lgr4/5dLKO n=4). **E.** HE staining revealed hepatic steatosis in a fraction of 6 months-old ND-fed Lgr4/5dLKO mice (arrowheads). **F.** 6 months-old HFD-fed Lgr4/5dLKO manifested lower serum t-Chol and c-HDL levels compared to WT mice under ND and HFD conditions, respectively (control ND n=9, Lgr4/5dLKO ND n=11, control HFD n=6, Lgr4/5dLKO HFD n=9; two way ANOVA, **** <0.0001, ** < 0.005). **G.** Elevated serum ALT and AST (n>5; t test, *= P < 0.05, **= P < 0.001) and t-BA (control n= 6, Lgr4/5dLKO n=9; t test, *= P < 0.05) levels in 6 months-old Lgr4/5dLKO compared to WT mice upon HFD. Results were expressed as ± SD. **H.** KRT19 staining in livers of 6 months-old Lgr4/5dLKO mice subjected to HFD showed that steatosis occurred in all zones Z1, Z2 and Z3 throughout the liver parenchyma (one section of moderate (a) and one of more pronounced (b) steatosis shown).

A similar but more pronounced phenotype was observed in a cohort of 6 months-old Lgr4/5dLKO mice. Histological analysis revealed that 60% of the ND-fed Lgr4/5dLKO female mice analyzed had developed spontaneous hepatic steatotic foci (Fig. 1E). Irrespective of sex and diet, HFD-fed Lgr4/5dLKO mice displayed reduced levels of serum t-Chol and c-HDL compared to control animals (Fig. 1F). Increased serum AST, ALT and t-BA levels were evident in HFD-fed Lgr4/5dLKO mice (Fig. 1G). Importantly, steatosis was not reflected by increased body weight in 6 months HFD-fed Lgr4/5dLKO mice (Fig. S1B). Liver steatosis occurred in all zones 1-3, with a higher preference for zones 1 and 2 over zone 3 in regions of moderate steatosis, as evidenced by monitoring the presence of lipid droplets in livers of 6 months-old HFD-fed Lgr4/5dLKO and WT mice relative to the biliary marker KRT19 (Fig. 1H).

### Lgr4/5dLKO mice revealed a NASH-like phenotype with fibrosis

To investigate whether the steatotic phenotype in 6 months-old HFD-fed LGR4/5dLKO mice was concomitant with other features of liver disease, we determined the presence of inflammatory cells. Steatosis was accompanied by hepatic inflammation, as evidenced by increased presence of Ionized Calcium-Binding Adapter Molecule 1 (IBA1) macrophages in Lgr4/5dLKO compared to WT mice (Fig. 2A). Furthermore, serum levels of the proinflammatory cytokines Interleukin 6 (IL6) and Tumor necrosis factor alpha (TNFα) were elevated in 6 months-old HFD-fed Lgr4/5dLKO (Fig. 2B). 40% of Lgr4/5dLKO animals developed porto-portal and porto-central bridging fibrosis upon HFD, whereas focal peri-cellular fibrosis and macro-vesicular steatosis were evident mainly along the fiber secta in the remaining 60% of animals (Fig. 2C). Increased Alpha Smooth Muscle Actin (αSMA) levels were present in livers of 6 months-old HFD-fed Lgr4/5dLKO mice (Fig. 2D), indicative of enhanced hepatic stellate cell (HSC) activation. To monitor the predisposition of Lgr4/5dLKO mice to liver fibrosis using a non-invasive diagnostic device, we subjected 3 and 6 months-old ND-fed Lgr4/5dLKO and control mice to liver ultrasound elastography. While liver stiffness in 3 months-old mice was comparable (data not shown), 6 months-old ND-fed Lgr4/5dLKO female mice tended to have stiffer livers compared to control animals (Fig. S1C). These findings are in line with increased fibrotic manifestation observed in 6 months-old HFD-fed Lgr4/5dLKO mice shown in Fig. 2C,D. In summary, our data strongly support the notion that mice with decreased Wnt/β-catenin signaling upon Lgr4/5dLKO deletion were prone to develop a NASH-like phenotype accompanied by fibrosis.

**Figure 2.**
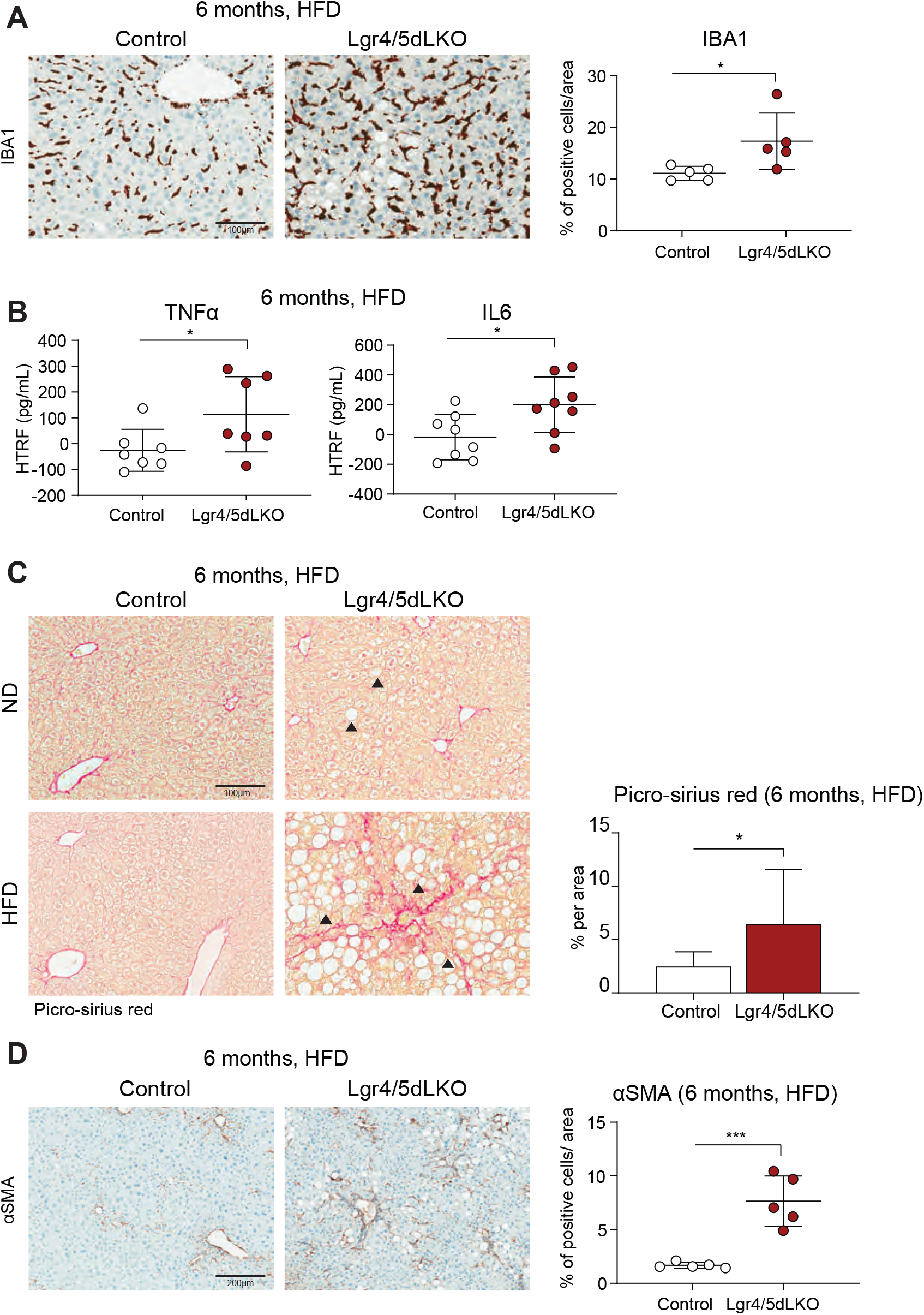
Liver inflammation with fibrosis in Lgr4/5dLKO mice. **A.** IBA1 staining revealed severe inflammation in livers of 6 months-old Lgr4/5dLKO mice upon HFD. Left, IBA1 staining; right, quantification of IBA1 staining (n>5; t test, *= P < 0.05); results were expressed as ± SD. **B.** Elevated serum levels of the proinflammatory cytokines IL6 and TNFα. Experimental animal conditions and quantification are described in Fig. 2A. **C.** Picro-sirius red staining on liver sections of 6 months-old ND-fed and HFD-fed (4 weeks) female mice. Spontaneous steatosis in ND-fed and macrovesicular steatosis in HFD-fed mice was detected in Lgr4/5dLKO mice (arrowheads) concomitant with increased collagen levels (picro-sirius red). Lgr4/5dLKO mice also developed bridging fibrosis upon HFD. Picro-sirius red quantification demonstrated that 6 months-old Lgr4/5dLKO mice subjected to HFD developed enhanced liver fibrosis (control n=10, Lgr4/5dLKO n= 9; *= P < 0.05). Results were expressed as ± SD. Experimental animal conditions as described in Fig. 1F. **D.** αSMA staining revealed increased HSC activation in 6-months old HFD-fed Lgr4/5dLKO compared to control animals (n=5, t test ***= P < 0.0001). Results were expressed as ± SD.

### Liver steatosis correlated with loss of BA secretion from Lgr4/5dLKO hepatocytes

Lipid accumulation can result from dysregulated free FA uptake from blood, enhanced *de novo* lipogenesis (DNL), increased chylomicron remnant hepatic uptake or diminished secretion [33]. To address the mechanism of steatosis in our Lgr4/5dLKO mice, we performed bulk RNAseq analysis on livers derived from 1 month-old Lgr4/5dLKO and control mice in order to capture causative transcriptome changes that preceded the phenotypic manifestation observed in 3 months-old mice. Livers of 1 month-old Lgr4/5dLKO mice did not display any visible lipid accumulation (Fig. S1D). Yet, the transcriptome analysis revealed several genes that were strongly down- or mildly upregulated (Fig. 3A,a). Among the former, we identified several genes encoding transporter proteins, indicative of altered intercellular transport function in livers of Lgr4/5dLKO mice. We performed pathway enrichment analysis on the down- and upregulated genes to gain mechanistic insights into the biological consequences of Lgr4/5 gene disruption. The main pathways downregulated comprised metabolism and synthesis of lipids, steroids, BA’s and bile salts as well as transport of bile salts, organic acids, small molecules and solute carrier (SLC)-mediated transmembrane transport (Fig. 3A,b). Upregulated pathways consisted of extracellular matrix organization and degradation, integrin and non-integrin interactions, collagen crosslinking, trimerization and degradation, glycosylation, and Neural Cell Adhesion Molecule (NCAM) and Platelet Derived Growth Factor (PDGF) signaling (Fig. 3A,c). In summary, our transcriptome data suggested a deficit in lipid and BA metabolism and an increased extracellular matrix remodeling propensity in the Lgr4/5dLKO hepatocytes, imposing compromised cellular function.

**Figure 3.**
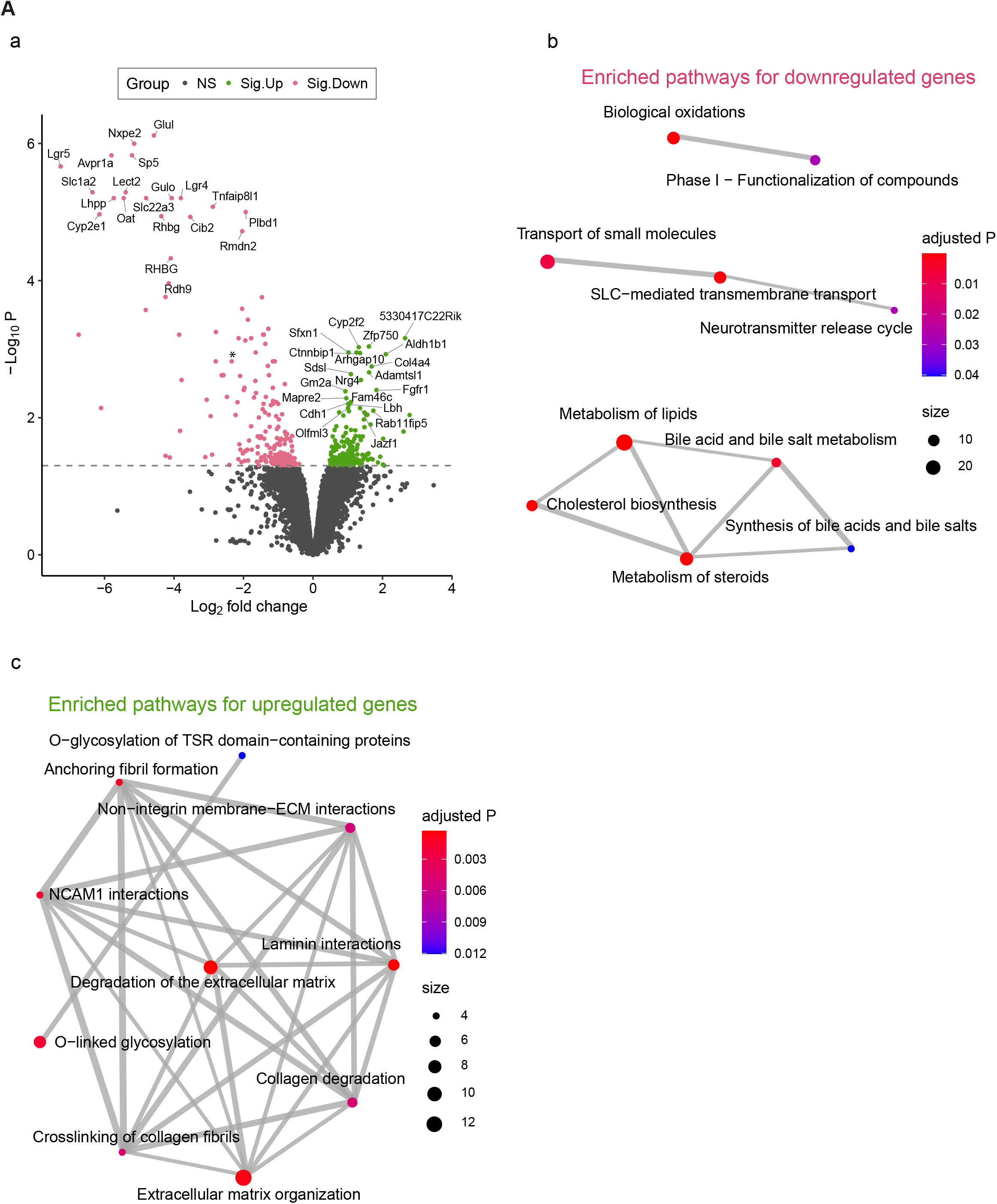
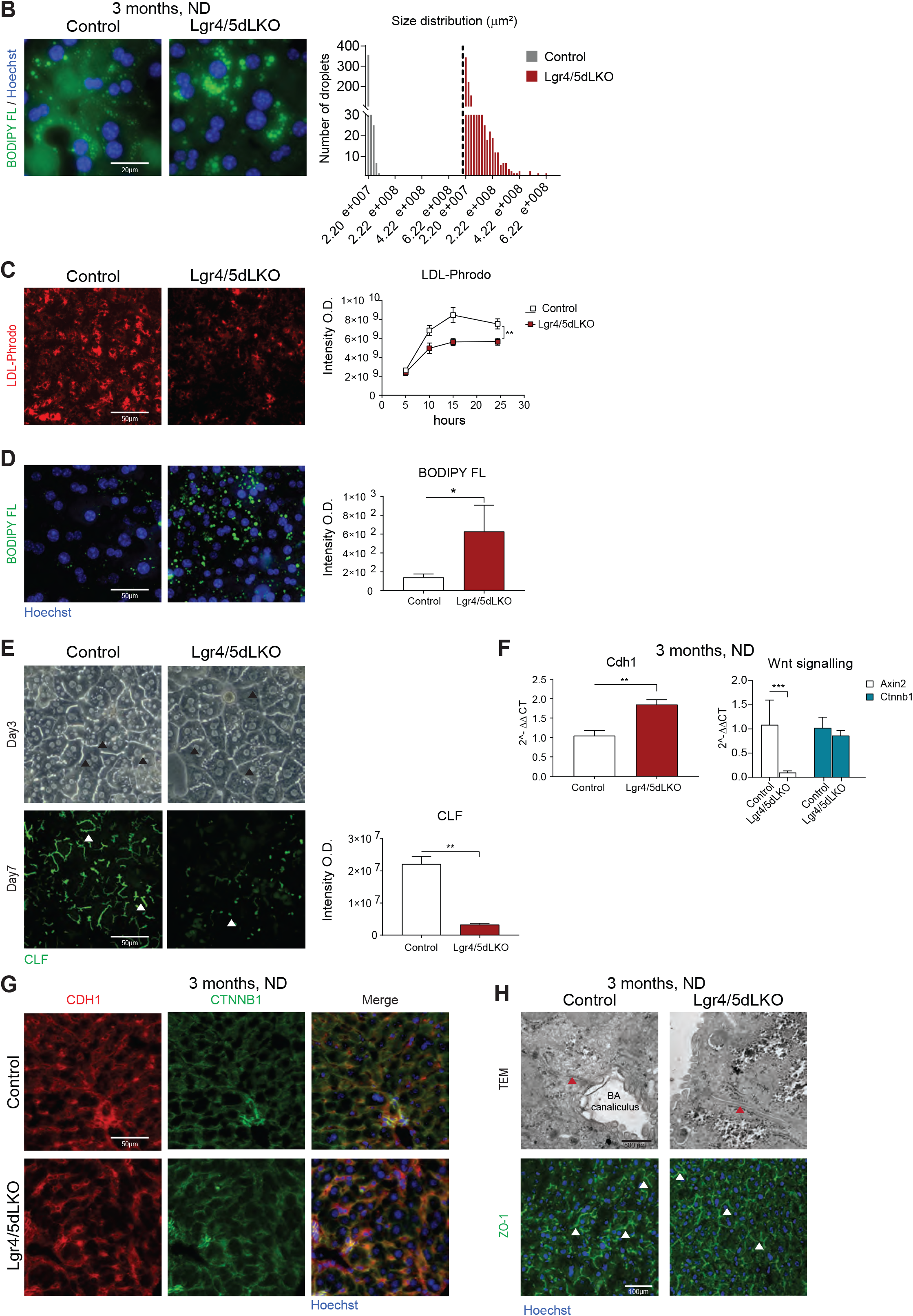
Hepatic lipid accumulation caused by decreased BA secretion in Lgr4/5dLKO hepatocytes. **A.** Livers of 1 month-old ND-fed Lgr4/5dLKO mice revealed a differential regulation of a subset of genes compared to WT control livers. (**a**) Volcano plot of differential gene expression analysis between Lgr4/5dLKO and control mice. P < 0.05 was used to define statistical significance. Only the top 20 up- or down-regulated genes are labelled with gene symbols. Significantly upregulated (Sig.Up) and downregulated (Sig.Down) genes in Lgr4/5dLKO mice are shown in green and pink, respectively. *, *Cldn2* expression decreased by −2.34 (log_2_-fold) in Lgr4/5dLKO compared to WT livers, adjusted p-value (P) 0.0015. (**b-c**) Pathway enrichment analysis for the 217 downregulated (**b**) and 203 upregulated (**c**) genes. Dot size represents the number of genes involved in each pathway. Dot color indicates the statistical significance (adjusted p-value) for each pathway. The thickness of the line connecting two pathways reflects the number of shared genes between the pathways. The more the shared genes, the thicker the line. **B.** BODIPY FL staining of primary hepatocytes isolated from 3 months-old ND-fed mice. Frequency distribution of lipid droplet-size (area expressed in μm^2^) was measured 24 hours after seeding. Lipid droplets in Lgr4/5dLKO hepatocytes differed in number and size from control cells (number of lipid droplets analyzed control > 400, Lgr4/5dLKO > 1500). **C.** Hepatocytes from 6 weeks-old Lgr4/5dLKO mice revealed significantly reduced LDL vesicle uptake compared to control cells (biological replicates n=2, technical replicates n=6; two-way ANOVA with Sidak’s test ****= P < 0.0001). LDL-Phrodo (red dots) images referred to the 24 h time point. **D.** BODIPY FL staining on hepatocytes at 24 h in culture after onset of LDL-Phrodo addition. Note the abundant content of endogenous lipid droplets in hepatocytes from Lgr4/5dLKO mice (biological replicates n=2, technical replicates controls n=6, Lgr4/5LdKO n=5; t test, *= P = 0.08). Results were expressed as ± SEM. **E.** Bright field and fluorescent images of collagen-matrigel 3D hepatocyte cultures 3 Days after cell seeding. The canaliculi network in Lgr4/5dLKO and control hepatocytes is exemplified by arrowheads. CLF assays demonstrated a loss of BA secretion from Lgr4/5dLKO hepatocytes on Day 7 (technical replicates n=3; t test, **= P < 0.001). Results were expressed as ± SEM. **F.** While *Axin2* gene expression was abolished in Lgr4/5dLKO mice, *Ctnnb1* gene expression was comparable to control animals (n=5; t-test ****= P < 0.0001). *Cdh1* gene expression in livers of Lgr4/5dLKO mice was slightly increased (n=6; t test, **= P < 0.001). Results were expressed as ± SD. **G.** CDH1 and CTNNB1 immunofluorescence staining on paraffin-embedded liver sections of 3 months-old ND-fed Lgr4/5dLKO and control animals showed that the co-localization of CDH1 and CTNNB1 was intact in the Lgr4/5dLKO mice. **H.** TEM and ZO-1 staining confirmed the presence of intact TJ’s in Lgr4/5dLKO mice (arrowheads).

In order to investigate how the reduced serum lipid levels in Lgr4/5dLKO mice correlated with hepatic lipid content, freshly isolated primary hepatocytes from 3 months-old ND-fed Lgr4/5dLKO and control mice were starved overnight in basal serum-free medium, and the lipid content was evaluated using a hydrophobic BODIPY FL dye. The *ex vivo* cultures confirmed that the Lgr4/5dLKO hepatocytes contained lipids that were stored in enlarged droplets compared to control hepatocytes (Fig. 3B). To assess whether lipid accumulation in Lgr4/5dLKO hepatocytes was caused by altered clearance of low density lipoprotein (LDL) from plasma, we tested the competence of Lgr4/5dLKO hepatocytes to take up LDL vesicles *ex vivo*. Freshly isolated hepatocytes from livers of both 6 weeks-old control and Lgr4/5dLKO mice were maintained as 2D monolayer cultures as depicted in Fig. S2A. On Day 2, hepatocytes were incubated with LDL vesicles conjugated to the red fluorophore Phrodo (LDL-Phrodo), and the kinetics of uptake was determined by measurements after 5, 10, 15, 20 and 24 hours of LDL-Phrodo addition. To distinguish between newly incorporated LDL vesicles and lipids existing in lipid droplets, LDL-Phrodo was removed after 24 hours (Day 3), and cells were incubated for 1 minute with BODIPY FL. Lgr4/5dLKO hepatocytes revealed a reduced LDL vesicle uptake compared to control cells (Fig. 3C) but contained abundant levels of endogenous lipid droplets (Fig. 3D). These results demonstrated that enhanced lipid accumulation in Lgr4/5dLKO hepatocytes was not a consequence of enhanced LDL uptake. In line with our findings on compromised lipid and BA metabolism in Lgr4/5dLKO mice, decreased expression of *Cldn2* suggested that the BA-mediated export mechanism was altered (Fig. 3A,a). To assess the BA secretory ability of Lgr4/5dLKO hepatocytes, primary hepatocytes were isolated from livers of 6 weeks-old ND-fed Lgr4/5dLKO and control mice and cultivated in a 3D collagen-matrigel sandwich format over 7 days, the period necessary to recreate functional canaliculi *ex vivo* [34]. The fluorescent BA analogue cholyl-*L*-lysyl-fluorescein (CLF) was utilized to quantify the BA efflux from hepatocytes according to the scheme depicted in Fig. S2B. Initiation of canalicular network formation in control and Lgr4/5dLKO hepatocytes started two days after seeding, whereby control cells showed an increased speed of canalicular structure development. On Day 3, both control and Lgr4/5dLKO hepatocytes cultures displayed similarly organized canalicular networks. On Day 7, Lgr4/5dLKO hepatocytes revealed strikingly reduced CLF secretion into BA canaliculi compared to control cells (Fig. 3E). In humans, defects in canalicular secretion with progressive intrahepatic trapping of BAs induce hepatocellular injury, biliary fibrosis and end stage liver disease [35]. To determine whether the compromised BA flow was due to altered hepatic junctions in Lgr4/5dLKO mice, we evaluated the integrity of adherens junctions by studying the hepatic localization of Cadherin 1 (CDH1) [36], whose expression was increased in livers of 3 months-old ND-fed Lgr4/5dLKO mice (Fig. 3F). CDH1 is known to interact with the Wnt pathway effector protein β-Catenin (CTNNB1), whose expression was unaltered, whereas the Wnt/β-catenin target gene Axis Inhibition Protein 2 (AXIN2) was strongly reduced (Fig. 3F). CDH1 and CTNNB1 revealed a similar co-localization in livers of 3 months-old ND-fed Lgr4/5dLKO and control animals, respectively, suggesting that the interaction of these proteins was unaltered in adherens junctions (Fig. 3G). Furthermore, TJ’s of 3 months-old ND-fed Lgr4/5dLKO mice appeared morphologically intact, as analyzed by transmission electron microscopy (TEM) and immunofluorescence analysis of the tight junction protein Zona Occludens Protein-1 (ZO-1) (Fig. 3H). Similarly, livers of *Cldn2^-/-^* mice revealed a normal bile canaliculi structure [8]. Our data implied that BA canaliculi functionality was not due to abnormal TJ formation. These results suggested that loss of BA secretion in Wnt/β-catenin-impaired mouse livers were caused by changes in cellular physiology and ECM remodeling, rather than by abnormal hepatic junctions.

### Livers of Lgr4/5dLKO mice revealed altered lipid profiles

Triggered by the steatotic liver phenotype in Lgr4/5dLKO mice, we performed lipid analysis on livers of 6 months-old Lgr4/5dLKO and control mice that were fed with either ND or HFD. HFD-fed Lgr4/5dLKO mice displayed an upregulation of several lipid species compared to ND, in particular triglycerols (TG). In contrast, fewer lipids were upregulated in control mice comparing HFD to ND conditions (Fig. 4A). Specifically, in HFD-fed Lgr4/5dLKO mice, we observed 45 different species of TG [37], 16 diacylglycerols (DGs), 10 oxidized triacylglycerols (OxTG) and 8 triacylglycerol estolides (TG-EST). Regarding the fatty acyl composition of significantly upregulated TGs, we observed that livers of Lgr4/5dLKO mice contained a more diverse FA pool including several polyunsaturated forms (*e.g.* 22:3, 22:4, 22:5) (Fig. 4B; square brackets denote polyunsaturated TGs). The complete list of increased and decreased lipid species upon HFD compared to ND, both in control and Lgr4/5dLKO mice, is shown in Table S1. We concluded that Lgr4/5dLKO mice exhibited a different lipid response to a HFD challenge compared to control mice, revealing altered lipid homeostasis.

**Figure 4:**
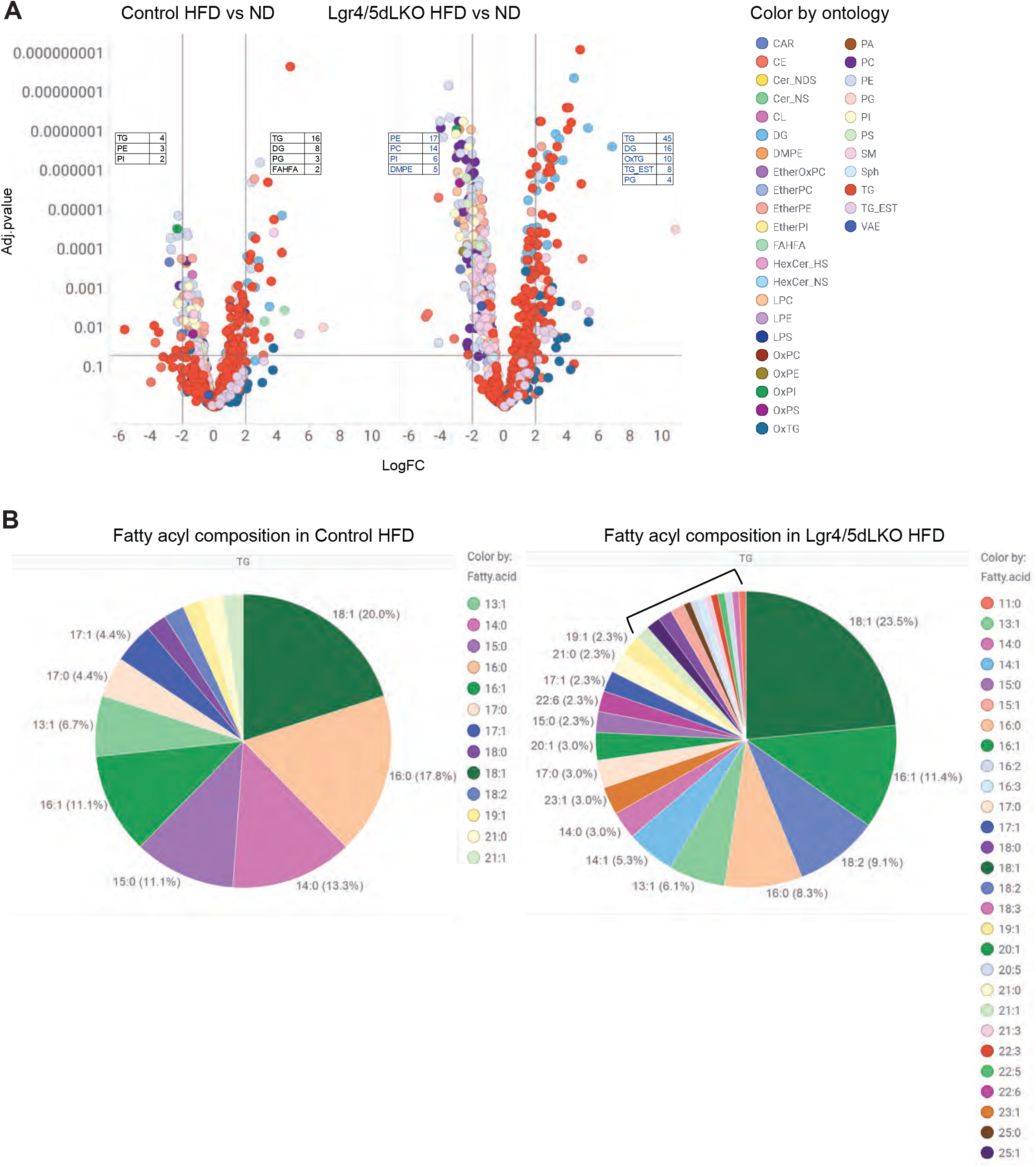
Lipid changes in livers of Lgr4/5dLKO mice. **A.** Volcano plots depicting lipid species found increased and decreased in livers of 6 months-old HFD-*versus* ND-fed Lgr4/5dLKO and control mice, respectively. HFD-fed Lgr4/5dLKO mice showed a significantly higher number of triacylglycerols (TG), diacylglycerols (DG), oxidized triacylglycerols (OxTG) and triacylglycerol estolides (TG-EST) compared to ND. Lipids changes with adj. p value > 0.05 and log_2_ fold change > 2 were considered as significantly changed. Tables depicted in volcano plots (black fonts for control, blue fonts for Lgr4/5dLKO) display the number of different lipid species identified, grouped by ontology, as either increased or decreased upon HFD compared to ND (right and left, respectively, in each graph). Abbreviations were used according to http://prime.psc.riken.jp/compms/msdial/lipidnomenclature.html. **B.** Pie chart representation of the distribution of specific TG fatty acyl species in HFD-fed Lgr4/5dLKO and control mice, respectively (expressed in %). Note that several fatty acyl species were only detected in livers of Lgr4/5dLKO mice, a fraction as polyunsaturated fatty acyl form (marked by bracket).

## Discussion

Increasing evidence suggests that Wnt/β-catenin signaling plays a role in both exogenous and endogenous lipid trafficking and biliary homeostasis, but the physiological consequences of Wnt/β-catenin pathway activity on hepatic metabolic functions and NAFLD development are incompletely understood. To address this question, we investigated phenotypic manifestations in liver and serum of mice devoid of Lgr4/5 in the liver, two components that are crucial for regulating liver Wnt/β-catenin pathway activity [19]. Our data showed that abrogation of Wnt/β-catenin activity led to fat accumulation in the liver, partly due to defective BA secretion, concomitant with increased liver inflammation and a predisposition to fibrosis.

The increased steatosis and resistance to diet-induced obesity that we observed in our Lgr4/5dLKO mice fed with HFD is in line with findings from β-catenin KO mice [21]. Likewise, *LDL-receptor related protein 6* (*Lrp6*) mutant mice showed enhanced hepatic *de novo* lipogenesis, lipid and cholesterol biosynthesis, resulting in elevated hepatic fat, triglycerol (TG) and cholesterol ester content, progressing towards NAFLD [24]. Loss-of-function point mutations in the *LRP6* gene lead to early onset of serum hyperlipidemia and metabolic syndrome in humans and engineered mice [24, 38]. Hepatocyte-specific β-catenin knockout mice developed hepatic steatosis when fed with methionine- and choline-deficient (MCD) diet but not with HFD. Collectively, this suggests that abrogated Wnt/β-catenin promotes NAFLD and NASH. In contrast, Wnt/β-catenin activation upon hepatocyte-specific ectopic β-catenin expression in mice fed with HFD or through loss of the Znrf3/Rnf43 module was described to cause hepatic steatosis [20, 21]. These studies provided insights about the role of the β-catenin transcriptional regulator in the context of liver metabolic disorders. It remains unclear why *Znrf3/Rnf43* deletion mediated by AAV8-Cre or Alb-Cre ERT2 caused NAFLD and NASH [20], whereas our mice with Ad5-Cre-mediated deletion of *Znrf3/Rnf43* showed the opposite phenotype. While our control mice developed significant age-related steatosis, aged *Znrf3/Rnf43* KO mice were protected from hepatic fat accumulation. We observed moderate steatosis in a subset of liver tumors that developed in our *Znrf3/Rnf43* KO mice, but the degree of steatosis was less than in livers of age-matched control littermates [22]. While our data supported the notion that abrogated Wnt/β-catenin promoted NALFD, more research is required by studying the detailed differences in the two Znrf3/Rnf43 KO models to explain this discrepancy. Yet, Wnt pathway effectors such as LRP5/6 receptors and β-catenin are involved in several physiological functions distinct from canonical Wnt/β-catenin transcriptional activation. For instance, LRP5/6 contain three copies of the LDL type-A motif, known to mediate the binding and the endocytosis of lipoproteins [24]. Similarly, β-catenin is a protein involved in regulation and coordination of cell-cell junctions, a structural cellular component whose integrity is coupled with TJ formation (reviewed in [39]). Therefore, it is plausible that the impact of LRP5/6 and β-catenin on NAFLD etiology results from a combination of impacted Wnt signaling, altered lipoprotein uptake and cell-cell contact.

NAFLD etiology can be the result of a combination of abnormalities in fat and amino acid metabolism, orchestrated by pericentral and periportal hepatocytes. Although the role of Wnt/β-catenin signaling in metabolic liver zonation is well characterized, the consequences of impaired metabolic process and how they may promote NAFLD are less clear. In NAFLD patients, lipid accumulation normally initiates in lipid droplets within pericentral hepatocytes [40, 41]. This lipid zonation is gradually lost during disease progression [42]. In contrast to Wnt/β-catenin KO mice which exhibited predominant periportal steatosis upon HFD feeding [21], we identified steatosis in a non-zonated manner. These results imply that hepatic lipid droplet accumulation in HFD-fed Lgr4/5dLKO mice appeared as a consequence, rather than a cause, of loss of metabolic liver zonation.

Wnt/β-catenin signaling was found to regulate BA biosynthesis in perivenous hepatocytes [6, 7]. Liver-specific deletion of the canonical Wnt pathway effector protein β-catenin leads to accumulation of hepatic cholesterol and BAs, elevation of serum bilirubin, increased steatohepatitis and fibrosis in mice fed with a steatogenic MCD diet [23], suggesting a putative defect in BA secretion due to dilated and tortuous bile canaliculi [43]. Inactivation of Wnt/β-catenin signaling caused strongly reduced expression of the perivenous membrane protein CLDN2, which participates in the formation of intercellular TJs. CLDN2 is involved in BA and water flow regulation in bile canaliculi, and deletion of *Cldn2* in polarized hepatocyte-like cells caused a disruption of functional bile canaliculus-like structures [8, 9]. Our *ex vivo* hepatocyte and transcriptomic data now extended these findings and implied that BA production and secretion were compromised in the Lgr4/5dLKO mice. Lgr4/5 deletion promoted loss of *Cldn2* gene expression, which was accompanied by reduced BA secretion from hepatocytes, suggesting a potential impairment of the endogenous lipid export pathway. It remains to be determined whether reduction of transmembrane transport mediated by SLCs identified in our transcriptome analysis in livers of the Lgr4/5dLKO mice affected cellular BA import or export.

The endogenous lipid trafficking route comprises the import and export of lipids into and from the liver by LDL and VLDL particles, respectively. Inadequate export of VLDL represents a central, incompletely understood mechanism in patients developing NAFLD. While individuals with simple steatosis exhibit increased VLDL-TG secretion, patients harboring genetic mutations that cause abnormal VLDL assembly or reduced VLDL export manifest an augmented risk to develop NASH [44, 45]. Wnt/β-catenin signaling was found to impinge on VLDL and the VLDL receptor (VLDLR). Knockdown of the VLDLR in retinal pigment epithelium resulted in the elevation of LRP6 protein levels and activation of Wnt/β-catenin signaling, whereas overexpression of VLDLR suppressed Wnt signaling [46]. Our Lgr4/5dLKO mice unveiled sub-physiological serum t-Chol and c-HDL but increased serum VLDL-cholesterol and VLDL-TG levels, likely due to a compensatory response towards intracellular hepatic lipid accumulation. Despite hepatic steatosis, these animals maintained normal body weight. Upon HFD challenge, Lgr4/5dLKO mice developed liver inflammation and fibrosis, and serum contained elevated AST and ALT levels.

Livers of Lgr4/5dLKO mice contained a different lipid profile compared to control mice, revealing that loss of liver zonation affected lipid homeostasis. Recent evidence has described LGR4 as a key regulator in lipid metabolism, both in adipose tissue and in liver [47, 48]. TG and their precursors DG were the predominant lipids that accumulated in livers of our 6 months-old HFD-fed Lgr4/5dLKO and control mice. We identified several TG comprising fatty acyl species in their polyunsaturated forms (*e.g.,* 22:1, 22:3 22:5, 22:6) in livers of Lgr4/5dLKO mice. Therefore, it is conceivable that the incapability of Lgr4/5dLKO hepatocytes to secrete BAs, in addition to defective hepatic FA oxidation by abrogated Wnt/β-catenin signaling [21], led to increased FA unsaturation and subsequent storage in large lipid droplets. Interestingly, we identified increased amounts of TG-ESTs in livers of 6 months-old HFD-fed Lgr4/5dLKO mice. TG-EST are TG-bound FA esters of hydroxy FAs (FAHFAs) and represent a major storage form of bioactive free FAHFAs [49]. FAHFAs represent a large family of lipids with poorly understood biological functions, including immunomodulatory activities [50]. It is plausible that the presence of these lipids reflected the need of the hepatocytes to cope with increased hepatic lipid load and/or to induce liver inflammation, causing the onset of NASH.

Our data revealed how loss of metabolic zonation upon liver-specific *Lgr4/5* ablation led to multiple molecular alterations that predisposed livers to steatosis and NAFLD progression. These events were manifested by transcriptional and lipid changes in the liver affecting not only hepatocyte physiology but also inflammation and HSC activation. Insights into this complex interaction network will likely result in the formation of new therapeutic hypotheses for the treatment of NAFLD.

## Abbreviations

Alb-Cre: Cre recombinase under the hepatocyte-specific albumin promoter
αSMA: Alpha Smooth Muscle Actin
ALT: alanine aminotransferase
AST: aspartate aminotransferase
AXIN2: Axis inhibition protein 2
BA: bile acid
CDH1: Cadherin 1
c-HDL: cholesterol-high density lipoprotein
CLDN2: Claudin-2
CLF: cholyl-L-lysyl-fluorescein
CTNNB1: Catenin beta 1
DG: diacylglycerol
DNL: *de novo* lipogenesis
FA: fatty acid
FAHFA: fatty acid esters of hydroxy fatty acids
FPLC: fast protein liquid chromatography-based sizeexclusion
GTT: glucose tolerance test
HFD: high fat diet
HSC: hepatic stellate cells
IBA1: Ionized Calcium-Binding Adapter Molecule 1
IHC: immunohistochemistry
IL6: Interleukin 6
ISH: *in situ* hybridization
ITT: insulin tolerance test
KO: knock-out
KRT19: Keratin 19
LGR: Leucine-Rich RepeatContaining G Protein-Coupled Receptor
Lgr4/5dLKO: Leucine-Rich Repeat-Containing G Protein-Coupled Receptor 4 and 5 double liver knock-out
LDL: low density lipoprotein
LRP6: LDL-Receptor Related Protein 6
MCD: methionine- and choline-deficient diet
NAFLD: nonalcoholic fatty liver disease
NASH: nonalcoholic steatohepatitis
ND: normal diet
OxTG: oxidized triglycerol
RNF43: Ring Finger Protein 43
RSPO: Roof Plate-Specific Spondin
SD, SD: standard deviation
SEM: standard error of the mean
SLC: solute carrier
SWE: shear wave elastography
t-BA: total bile acids
t-Chol: total cholesterol
TEM: Transmission electron microscopy
TG: triglycerol
TG-EST: triacylglycerol estolides
TJ: tight junctions
TNFα: tumor necrosis factor alpha
VLDL: very low density lipoprotein
VLDLR: very low density lipoprotein receptor
Wnt: Wingless-related integration site
WT: wild-type
ZNRF3: Zinc And Ring Finger 3
ZO-1: Zona Occludens Protein-1

## Financial support

This study was supported by the Novartis Institutes for BioMedical Research Postdoctoral Program.

## Conflict of interest

All authors except M.L.M.G., A.G.M., C.G. and L.T. are or were employees of Novartis Pharma AG.

## Author’s contributions

E.S. and H.R. have conceived this study. E.S., C.P., M.L.M.G., V.B., B.F., P.OD., J.T., A.G.M., L.B., G.A., W.C., V.O., S.A., M.O., N.B., C.S., A.A. and S.L. have carried out the experiments. E.S., H.R., T.B., J.T., C.P., M.B., S.R., I.K., F.N., L.T., Z.Y.W. discussed and interpreted results. A.O., M.B., G.R., P.T., N.B., C.S., I.K., C.G. and J.T. provided the key materials and instructions for usage. E.S., J.T. and H.R. wrote the manuscript. H.R. and T.B. supervised the project.

## Acknowledgements

We thank Ludovic Perrot, Sabrina Silvia Surber, Nathalie Stuber, Martin van de Velde, Gabi Schutzius, Adrian Salathe, Berangere Gapp, Benjamin Kueng and Dominic Trojer for technical assistance. For helpful discussion and critical reading of the manuscript, we thank Klaus Seuwen, Susan Kirkland, Gabriele Hintzen, Felix Lohmann and Anne Granger.

---

**Figure S1. Phenotypic manifestations in Lgr4/5dLKO mice. A.** GTT and ITT performed on 3 months-old ND-fed mice (n=6). Results were expressed as ± SD. **B.** Body weight gain in 6 months-old Lgr4/5dLKO mice subjected to HFD compared to control animals (control n=10, Lgr4/5dLKO n=9). **C.** Ultrasound elastography was performed on 6 months-old ND-fed female mice. Lgr4/5dLKO mice tended to develop stiffer livers compared to control animals (control n=4, Lgr4/5dLKO n=4). The liver elasticity (kPa) was measured by shear wave elastography (SWE). **D.** HE staining on livers of 1 month-old ND-fed Lgr4/5dLKO and control animals. No visible anomalies were apparent in livers of Lgr4/5dLKO mice.

**Figure S2. *Ex vivo* primary hepatocyte culture protocols. A.** Schematic representation of protocol for isolation of primary hepatocytes followed by LDL-Phrodo uptake kinetic measurement and BODIPY FL staining. **B.** Schematic representation of protocol for isolation of primary hepatocytes followed by measurement of CLF secretion.

**Table S1. Lipid species found perturbed in 6 months-old HFD-compared to ND-fed Lgr4/5dLKO and control animals, respectively.** Decreased and increased lipids depicted on the left or right side of each volcano plot, respectively, in Fig. 4A. Adj. p value < 0.05, Log FC < −2 and > +2. Additionally, the adduct form (*e.g.* [M+H]) and the retention time (*e.g.* 1.078) for a given lipid were concatenated to the lipid name in order to handle double identifications.

## Notes

### Competing Interest Statement

All authors except M.L.M.G, A.G.M, C.G. and L.T. are or have been employed by and/or shareholders of Novartis Pharma AG.

## References

1. Ben-Moshe, S. and S. Itzkovitz, Spatial heterogeneity in the mammalian liver. Nat Rev Gastroenterol Hepatol, 2019. 16(7): p. 395–410.

2. Gebhardt, R., Metabolic zonation of the liver: regulation and implications for liver function. Pharmacol Ther, 1992. 53(3): p. 275–354.

3. Peng, K., Z. Mo, and G. Tian, Serum Lipid Abnormalities and Nonalcoholic Fatty Liver Disease in Adult Males. Am J Med Sci, 2017. 353(3): p. 236–241.

4. Bashiri, A., et al., Emerging role of cellular cholesterol in the pathogenesis of nonalcoholic fatty liver disease. Current Opinion in Lipidology, 2013. 24(3): p. 275–276.

5. Musso, G., R. Gambino, and M. Cassader, Cholesterol metabolism and the pathogenesis of non-alcoholic steatohepatitis. Progress in Lipid Research, 2013. 52(1): p. 175–191.

6. Behari, J., The Wnt/beta-catenin signaling pathway in liver biology and disease. Expert Rev Gastroenterol Hepatol, 2010. 4(6): p. 745–56.

7. Lemberger, U.J., et al., Hepatocyte specific expression of an oncogenic variant of beta-catenin results in cholestatic liver disease. Oncotarget, 2016. 7(52): p. 86985–86998.

8. Matsumoto, K., et al., Claudin 2 Deficiency Reduces Bile Flow and Increases Susceptibility to Cholesterol Gallstone Disease in Mice. Gastroenterology, 2014. 147(5): p. 1134-+.

9. Son, S., et al., Knockdown of tight junction protein claudin-2 prevents bile canalicular formation in WIF-B9 cells. Histochem Cell Biol, 2009. 131(3): p. 411–24.

10. MacDonald, B.T., K. Tamai, and X. He, Wnt/beta-catenin signaling: components, mechanisms, and diseases. Dev Cell, 2009. 17(1): p. 9–26.

11. Russell, J.O. and S.P. Monga, Wnt/beta-Catenin Signaling in Liver Development, Homeostasis, and Pathobiology. Annu Rev Pathol, 2018. 13: p. 351–378.

12. Benhamouche, S., et al., Apc tumor suppressor gene is the “zonation-keeper” of mouse liver. Dev Cell, 2006. 10(6): p. 759–70.

13. Ma, R., et al., Metabolic and non-metabolic liver zonation is established non-synchronously and requires sinusoidal Wnts. Elife, 2020. 9.

14. Yang, J., et al., beta-catenin signaling in murine liver zonation and regeneration: a Wnt-Wnt situation! Hepatology, 2014. 60(3): p. 964–76.

15. de Lau, W., et al., Lgr5 homologues associate with Wnt receptors and mediate R-spondin signalling. Nature, 2011. 476(7360): p. 293–7.

16. Hao, H.X., et al., ZNRF3 promotes Wnt receptor turnover in an R-spondin-sensitive manner. Nature, 2012. 485(7397): p. 195–200.

17. Koo, B.K., et al., Tumour suppressor RNF43 is a stem-cell E3 ligase that induces endocytosis of Wnt receptors. Nature, 2012. 488(7413): p. 665–9.

18. Ruffner, H., et al., R-Spondin potentiates Wnt/beta-catenin signaling through orphan receptors LGR4 and LGR5. PLoS One, 2012. 7(7): p. e40976.

19. Planas-Paz, L., et al., Corrigendum: The RSPO-LGR4/5-ZNRF3/RNF43 module controls liver zonation and size. Nat Cell Biol, 2016. 18(11): p. 1260.

20. Mastrogiovanni, G., et al., Loss of RNF43/ZNRF3 predisposes to Hepatocellular carcinoma by impairing liver regeneration and altering liver fat metabolism. bioRxiv, 2020: p. 2020.09.25.313205.

21. Behari, J., et al., beta-Catenin Links Hepatic Metabolic Zonation with Lipid Metabolism and Diet-Induced Obesity in Mice. American Journal of Pathology, 2014. 184(12): p. 3284–3298.

22. Sun, T., et al., ZNRF3 and RNF43 cooperate to safeguard metabolic liver zonation and hepatocyte proliferation. Cell Stem Cell, 2021.28(10): p. 1822–1837 e10.

23. Behari, J., et al., Liver-Specific beta-Catenin Knockout Mice Exhibit Defective Bile Acid and Cholesterol Homeostasis and Increased Susceptibility to Diet-Induced Steatohepatitis. American Journal of Pathology, 2010. 176(2): p. 744–753.

24. Go, G.W., et al., The Combined Hyperlipidemia Caused by Impaired Wnt-LRP6 Signaling Is Reversed by Wnt3a Rescue. Cell Metabolism, 2014. 19(2): p. 209–220.

25. Yu, G., et al., clusterProfiler: an R package for comparing biological themes among gene clusters. OMICS, 2012. 16(5): p. 284–7.

26. Yu, G. and Q.Y. He, ReactomePA: an R/Bioconductor package for reactome pathway analysis and visualization. Mol Biosyst, 2016. 12(2): p. 477–9.

27. McGuinness, O.P., et al., NIH experiment in centralized mouse phenotyping: the Vanderbilt experience and recommendations for evaluating glucose homeostasis in the mouse. Am J Physiol Endocrinol Metab, 2009. 297(4): p. E849–55.

28. Tsugawa, H., et al., A lipidome atlas in MS-DIAL 4. Nat Biotechnol, 2020. 38(10): p. 1159–1163.

29. De Livera, A.M., et al., Statistical methods for handling unwanted variation in metabolomics data. Anal Chem, 2015. 87(7): p. 3606–15.

30. Gapp, B., et al., Farnesoid X Receptor Agonism, Acetyl-Coenzyme A Carboxylase Inhibition, and Back Translation of Clinically Observed Endpoints of De Novo Lipogenesis in a Murine NASH Model. Hepatology Communications, 2020. 4(1): p. 109–125.

31. Chatrath, H., R. Vuppalanchi, and N. Chalasani, Dyslipidemia in patients with nonalcoholic fatty liver disease. Semin Liver Dis, 2012. 32(1): p. 22–9.

32. Fabbrini, E., et al., Alterations in adipose tissue and hepatic lipid kinetics in obese men and women with nonalcoholic fatty liver disease. Gastroenterology, 2008. 134(2): p. 424–431.

33. Mashek, D.G., Hepatic fatty acid trafficking: multiple forks in the road. Adv Nutr, 2013. 4(6): p. 697–710.

34. Germano, D., et al., Determination of liver specific toxicities in rat hepatocytes by high content imaging during 2-week multiple treatment. Toxicol In Vitro, 2015. 30(1 Pt A): p. 79–94.

35. Fickert, P. and M. Wagner, Biliary bile acids in hepatobiliary injury - What is the link? Journal of Hepatology, 2017. 67(3): p. 619–631.

36. Pradhan-Sundd, T. and S.P. Monga, Blood-Bile Barrier: Morphology, Regulation, and Pathophysiology. Gene Expr, 2019. 19(2): p. 69–87.

37. Alves-Bezerra, M. and D.E. Cohen, Triglyceride Metabolism in the Liver. Compr Physiol, 2017. 8(1): p. 1–8.

38. Mani, A., et al., LRP6 mutation in a family with early coronary disease and metabolic risk factors. Science, 2007. 315(5816): p. 1278–82.

39. Campbell, H.K., J.L. Maiers, and K.A. DeMali, Interplay between tight junctions & adherens junctions. Experimental Cell Research, 2017. 358(1): p. 39–44.

40. Chalasani, N., et al., Relationship of steatosis grade and zonal location to histological features of steatohepatitis in adult patients with non-alcoholic fatty liver disease. Journal of Hepatology, 2008. 48(5): p. 829–834.

41. Straub, B.K., et al., Differential pattern of lipid droplet-associated proteins and de novo perilipin expression in hepatocyte steatogenesis. Hepatology, 2008. 47(6): p. 1936–1946.

42. Hall, Z., et al., Lipid Zonation and Phospholipid Remodeling in Nonalcoholic Fatty Liver Disease. Hepatology, 2017. 65(4): p. 1165–1180.

43. Yeh, T.H., et al., Liver-Specific beta-Catenin Knockout Mice Have Bile Canalicular Abnormalities, Bile Secretory Defect, and Intrahepatic Cholestasis. Hepatology, 2010. 52(4): p. 1410–1419.

44. Dongiovanni, P., et al., Transmembrane 6 Superfamily Member 2 Gene Variant Disentangles Nonalcoholic Steatohepatitis From Cardiovascular Disease. Hepatology, 2015. 61(2): p. 506–514.

45. Fabbrini, E. and F. Magkos, Hepatic Steatosis as a Marker of Metabolic Dysfunction. Nutrients, 2015. 7(6): p. 4995–5019.

46. Lee, K., et al., Receptor heterodimerization as a novel mechanism for the regulation of Wnt/beta-catenin signaling. Journal of Cell Science, 2014. 127(22): p. 4857–4869.

47. Wang, J., et al., Ablation of LGR4 promotes energy expenditure by driving white-to-brown fat switch. Nat Cell Biol, 2013. 15(12): p. 1455–63.

48. Wang, F., et al., LGR4 acts as a link between the peripheral circadian clock and lipid metabolism in liver. Journal of Molecular Endocrinology, 2014. 52(2): p. 133–143.

49. Brejchova, K., et al., Distinct roles of adipose triglyceride lipase and hormone-sensitive lipase in the catabolism of triacylglycerol estolides. Proc Natl Acad Sci U S A, 2021. 118(2).

50. Riecan, M., et al., Branched and linear fatty acid esters of hydroxy fatty acids (FAHFA) relevant to human health. Pharmacol Ther, 2021: p. 107972.

