## Supplementary figures and images for "Loss of hepatic Lgr4 and Lgr5 promotes nonalcoholic fatty liver disease"

### Supplemental Figures 1-2

#### Figure S1

A

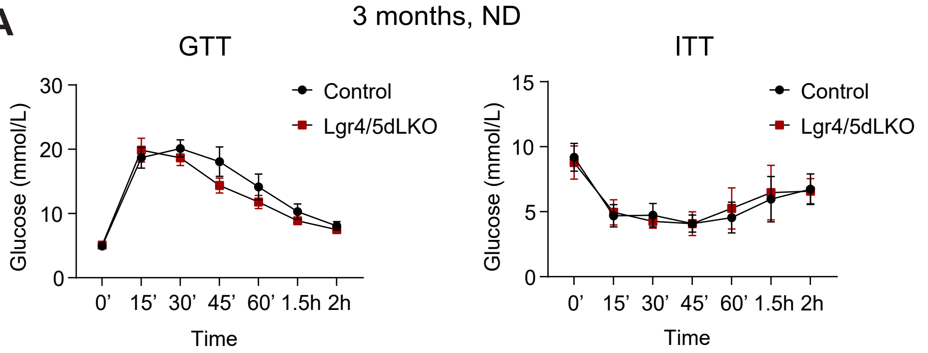


B

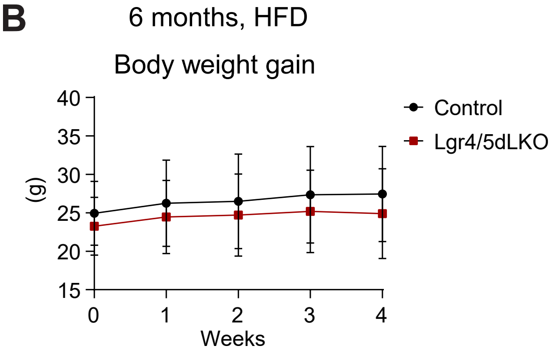


C

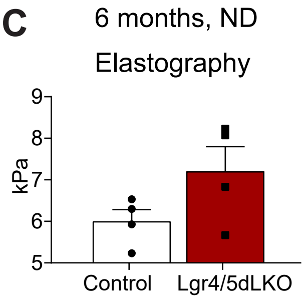


D

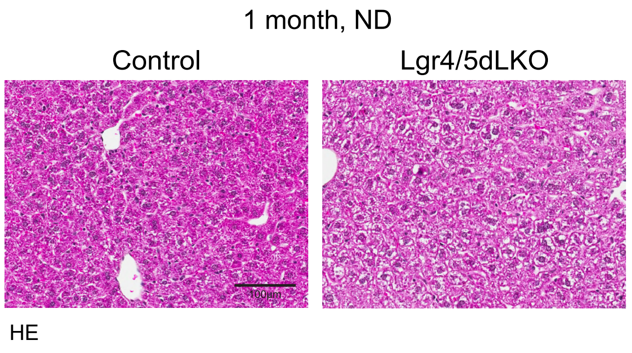

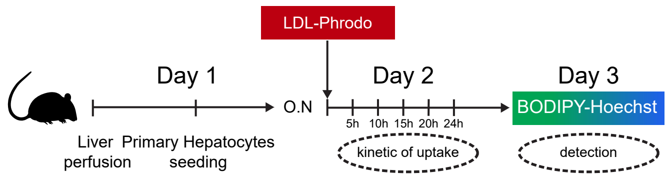

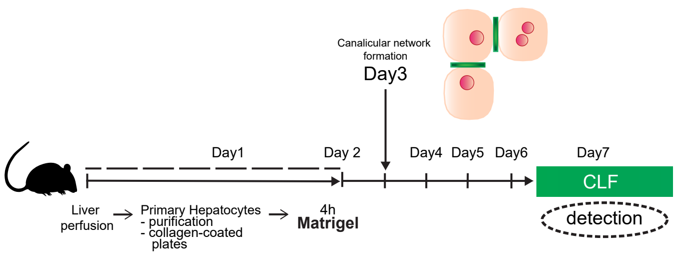
