## Supplemental Table 1 for "Loss of hepatic Lgr4 and Lgr5 promotes nonalcoholic fatty liver disease"

### Table S1

| Lipid name | Contrasts | LogFC | Adj.pvals | Ontology | Saturation | Number of Carbons |
|---|---|---|---|---|---|---|
| TG 64:8\|TG 18:1_18:1_28:6_[M+NH4]+_25.324 | Control HFD - Control ND | -5.676564933 | 0.010670344 | TG | Polyunsaturated | 64 |
| TG 64:8\|TG 18:1_18:1_28:6_[M+NH4]+_25.324 | Control HFD - Control ND | -5.676564933 | 0.010670344 | TG | Polyunsaturated | 64 |
| TG 62:14\|TG 18:2_22:6_22:6_[M+NH4]+_22.264 | Control HFD - Control ND | -3.792271253 | 0.008460391 | TG | Polyunsaturated | 62 |
| TG 62:14\|TG 18:2_22:6_22:6_[M+NH4]+_22.264 | Control HFD - Control ND | -3.792271253 | 0.008460391 | TG | Polyunsaturated | 62 |
| CE 24:5_[M+NH4]+_25.693 | Control HFD - Control ND | -3.73456656 | 0.022711149 | CE | Polyunsaturated | 24 |
| CE 24:5_[M+NH4]+_25.693 | Control HFD - Control ND | -3.73456656 | 0.022711149 | CE | Polyunsaturated | 24 |
| TG 60:13\|TG 16:1_22:6_22:6_[M+NH4]+_22.056 | Control HFD - Control ND | -3.531449854 | 0.01230322 | TG | Polyunsaturated | 60 |
| TG 60:13\|TG 16:1_22:6_22:6_[M+NH4]+_22.056 | Control HFD - Control ND | -3.531449854 | 0.01230322 | TG | Polyunsaturated | 60 |
| CAR 4:0_[M+H]+_1.078 | Control HFD - Control ND | -2.838085786 | 0.000204346 | CAR | Saturated | 4 |
| CAR 4:0_[M+H]+_1.078 | Control HFD - Control ND | -2.838085786 | 0.000204346 | CAR | Saturated | 4 |
| PE 36:6\|PE 16:1_20:5_[M-H]-_13.782 | Control HFD - Control ND | -2.764556555 | 5.05351E-05 | PE | Polyunsaturated | 36 |
| PE 36:6\|PE 16:1_20:5_[M-H]-_13.782 | Control HFD - Control ND | -2.764556555 | 5.05351E-05 | PE | Polyunsaturated | 36 |
| PE 36:6_[M+H]+_13.792 | Control HFD - Control ND | -2.744141126 | 9.58141E-05 | PE | Polyunsaturated | 36 |
| PE 36:6_[M+H]+_13.792 | Control HFD - Control ND | -2.744141126 | 9.58141E-05 | PE | Polyunsaturated | 36 |
| TG 62:13\|TG 18:1_22:6_22:6_[M+NH4]+_22.85 | Control HFD - Control ND | -2.430174542 | 0.036478435 | TG | Polyunsaturated | 62 |

| | | | | | | |
|---|---|---|---|---|---|---|
| TG 62:13\|TG 18:1_22:6_22:6_[M+NH4]+_22.85 | Control HFD - Control ND | -2.430174542 | 0.036478435 | TG | Polyunsaturated | 62 |
| PI 32:1;3O\|PI 16:0_16:1;3O_[M-H]-_13.687 | Control HFD - Control ND | -2.342315211 | 2.92661E-05 | OxPI | Monounsaturated | 32 |
| PI 32:1;3O\|PI 16:0_16:1;3O_[M-H]-_13.687 | Control HFD - Control ND | -2.342315211 | 2.92661E-05 | OxPI | Monounsaturated | 32 |
| PE 32:2\|PE 16:1_16:1_[M-H]-_14.665 | Control HFD - Control ND | -2.314969939 | 0.006536341 | PE | Polyunsaturated | 32 |
| PE 32:2\|PE 16:1_16:1_[M-H]-_14.665 | Control HFD - Control ND | -2.314969939 | 0.006536341 | PE | Polyunsaturated | 32 |
| PE 36:5_[M+H]+_15.288 | Control HFD - Control ND | -2.306013673 | 4.45278E-05 | PE | Polyunsaturated | 36 |
| PE 36:5_[M+H]+_15.288 | Control HFD - Control ND | -2.306013673 | 4.45278E-05 | PE | Polyunsaturated | 36 |
| PI 34:2\|PI 16:0_18:2_[M-H]-_14.837 | Control HFD - Control ND | -2.295380232 | 0.002199245 | PI | Polyunsaturated | 34 |
| PI 34:2\|PI 16:0_18:2_[M-H]-_14.837 | Control HFD - Control ND | -2.295380232 | 0.002199245 | PI | Polyunsaturated | 34 |
| PE 36:5\|PE 16:0_20:5_[M-H]-_15.295 | Control HFD - Control ND | -2.287587283 | 1.36717E-05 | PE | Polyunsaturated | 36 |
| PE 36:5\|PE 16:0_20:5_[M-H]-_15.295 | Control HFD - Control ND | -2.287587283 | 1.36717E-05 | PE | Polyunsaturated | 36 |
| PI 36:3\|PI 16:0_20:3_[M-H]-_15.261 | Control HFD - Control ND | -2.251810668 | 0.002322104 | PI | Polyunsaturated | 36 |
| PI 36:3\|PI 16:0_20:3_[M-H]-_15.261 | Control HFD - Control ND | -2.251810668 | 0.002322104 | PI | Polyunsaturated | 36 |
| PC 36:6\|PC 16:1_20:5_[M+HCOO]-_13.54 | Control HFD - Control ND | -2.187207727 | 0.000798658 | PC | Polyunsaturated | 36 |
| PC 36:6\|PC 16:1_20:5_[M+HCOO]-_13.54 | Control HFD - Control ND | -2.187207727 | 0.000798658 | PC | Polyunsaturated | 36 |
| PE O-40:9\|PE O-18:3_22:6_[M-H]-_15.593 | Control HFD - Control ND | -2.057651367 | 0.000163915 | EtherPE | Polyunsaturated | |

| | | | | | | |
|---|---|---|---|---|---|---|
| DG 36:1\|DG 18:0_18:1_[M+NH4]+_20.457 | Control HFD - Control ND | 2.038773742 | 0.011633935 | DG | Monounsaturated | 36 |
| DG 36:1\|DG 18:0_18:1_[M+NH4]+_20.457 | Control HFD - Control ND | 2.038773742 | 0.011633935 | DG | Monounsaturated | 36 |
| DG 33:1\|DG 15:0_18:1_[M+NH4]+_18.816 | Control HFD - Control ND | 2.073399704 | 0.000143891 | DG | Monounsaturated | 33 |
| DG 33:1\|DG 15:0_18:1_[M+NH4]+_18.816 | Control HFD - Control ND | 2.073399704 | 0.000143891 | DG | Monounsaturated | 33 |
| TG 47:2\|TG 14:0_16:1_17:1_[M+NH4]+_23.374 | Control HFD - Control ND | 2.073706333 | 0.012603154 | TG | Polyunsaturated | 47 |
| TG 47:2\|TG 14:0_16:1_17:1_[M+NH4]+_23.374 | Control HFD - Control ND | 2.073706333 | 0.012603154 | TG | Polyunsaturated | 47 |
| TG 47:2\|TG 14:0_16:1_17:1_[M+NH4]+_23.335 | Control HFD - Control ND | 2.083391995 | 0.012693892 | TG | Polyunsaturated | 47 |
| TG 47:2\|TG 14:0_16:1_17:1_[M+NH4]+_23.335 | Control HFD - Control ND | 2.083391995 | 0.012693892 | TG | Polyunsaturated | 47 |
| DG 38:2\|DG 18:1_20:1_[M+NH4]+_20.42 | Control HFD - Control ND | 2.102536689 | 0.000675956 | DG | Polyunsaturated | 38 |
| DG 38:2\|DG 18:1_20:1_[M+NH4]+_20.42 | Control HFD - Control ND | 2.102536689 | 0.000675956 | DG | Polyunsaturated | 38 |
| PG 35:2\|PG 17:1_18:1_[M-H]-_14.826 | Control HFD - Control ND | 2.151834528 | 0.006731401 | PG | Polyunsaturated | 35 |
| PG 35:2\|PG 17:1_18:1_[M-H]-_14.826 | Control HFD - Control ND | 2.151834528 | 0.006731401 | PG | Polyunsaturated | 35 |
| TG 47:2\|TG 14:0_15:0_18:2_[M+NH4]+_23.439 | Control HFD - Control ND | 2.185781697 | 0.005469803 | TG | Polyunsaturated | 47 |
| TG 47:2\|TG 14:0_15:0_18:2_[M+NH4]+_23.439 | Control HFD - Control ND | 2.185781697 | 0.005469803 | TG | Polyunsaturated | 47 |
| TG 55:2\|TG 16:0_18:1_21:1_[M+NH4]+_25.735 | Control HFD - Control ND | 2.237706713 | 0.000174727 | TG | Polyunsaturated | 55 |
| TG 55:2\|TG 16:0_18:1_21:1_[M+NH4]+_25.735 | Control HFD - Control ND | 2.237706713 | 0.000174727 | TG | Polyunsaturated | 55 |
| PG 39:4\|PG 18:1_21:3_[M-H]-_15.644 | Control HFD - Control ND | 2.259090236 | 0.012204564 | PG | Polyunsaturated | 39 |
| PG 39:4\|PG 18:1_21:3_[M-H]-_15.644 | Control HFD - Control ND | 2.259090236 | 0.012204564 | PG | Polyunsaturated | 39 |
| TG 47:3\|TG 13:1_16:1_18:1_[M+NH4]+_22.949 | Control HFD - Control ND | 2.292241737 | 0.000129021 | TG | Polyunsaturated | 47 |

| | | | | | | |
|---|---|---|---|---|---|---|
| TG 47:3\|TG 13:1_16:1_18:1_[M+NH4]+_22.949 | Control HFD - Control ND | 2.292241737 | 0.000129021 | TG | Polyunsaturated | 47 |
| PE 39:6_[M+H]+_16.363 | Control HFD - Control ND | 2.3403596 | 0.00029685 | PE | Polyunsaturated | 39 |
| PE 39:6_[M+H]+_16.363 | Control HFD - Control ND | 2.3403596 | 0.00029685 | PE | Polyunsaturated | 39 |
| PE 40:7_[M+H]+_16.288 | Control HFD - Control ND | 2.357739769 | 1.89031E-06 | PE | Polyunsaturated | 40 |
| PE 40:7_[M+H]+_16.288 | Control HFD - Control ND | 2.357739769 | 1.89031E-06 | PE | Polyunsaturated | 40 |
| TG 51:1\|TG 16:0_17:0_18:1_[M+NH4]+_25.313 | Control HFD - Control ND | 2.384488357 | 0.000874311 | TG | Monounsaturated | 51 |
| TG 51:1\|TG 16:0_17:0_18:1_[M+NH4]+_25.313 | Control HFD - Control ND | 2.384488357 | 0.000874311 | TG | Monounsaturated | 51 |
| DG 35:2\|DG 17:1_18:1_[M+NH4]+_18.874 | Control HFD - Control ND | 2.431535961 | 4.62426E-05 | DG | Polyunsaturated | 35 |
| DG 35:2\|DG 17:1_18:1_[M+NH4]+_18.874 | Control HFD - Control ND | 2.431535961 | 4.62426E-05 | DG | Polyunsaturated | 35 |
| PE O-37:5\|PE O-17:1_20:4_[M-H]-_17.301 | Control HFD - Control ND | 2.531078219 | 1.53213E-06 | EtherPE | Polyunsaturated | |
| PE O-37:5\|PE O-17:1_20:4_[M-H]-_17.301 | Control HFD - Control ND | 2.531078219 | 1.53213E-06 | EtherPE | Polyunsaturated | |
| TG 45:1\|TG 14:0_15:0_16:1_[M+NH4]+_23.329 | Control HFD - Control ND | 2.556271035 | 0.012205143 | TG | Monounsaturated | 45 |
| TG 45:1\|TG 14:0_15:0_16:1_[M+NH4]+_23.329 | Control HFD - Control ND | 2.556271035 | 0.012205143 | TG | Monounsaturated | 45 |
| TG 55:1\|TG 16:0_21:0_18:1_[M+NH4]+_26.172 | Control HFD - Control ND | 2.558170281 | 0.018989618 | TG | Monounsaturated | 55 |
| TG 55:1\|TG 16:0_21:0_18:1_[M+NH4]+_26.172 | Control HFD - Control ND | 2.558170281 | 0.018989618 | TG | Monounsaturated | 55 |
| DG 36:2\|DG 18:1_18:1_[M+NH4]+_19.43 | Control HFD - Control ND | 2.589351262 | 0.000898171 | DG | Polyunsaturated | 36 |
| DG 36:2\|DG 18:1_18:1_[M+NH4]+_19.43 | Control HFD - Control ND | 2.589351262 | 0.000898171 | DG | Polyunsaturated | 36 |
| TG 47:2\|TG 16:0_13:1_18:1_[M+NH4]+_23.634 | Control HFD - Control ND | 2.62977506 | 5.13937E-05 | TG | Polyunsaturated | 47 |
| TG 47:2\|TG 16:0_13:1_18:1_[M+NH4]+_23.634 | Control HFD - Control ND | 2.62977506 | 5.13937E-05 | TG | Polyunsaturated | 47 |
| DG 37:2\|DG 18:1_19:1_[M+NH4]+_19.945 | Control HFD - Control ND | 2.776285482 | 0.000187179 | DG | Polyunsaturated | 37 |

| | | | | | | |
|---|---|---|---|---|---|---|
| DG 37:2\|DG 18:1_19:1_[M+NH4]+_19.945 | Control HFD - Control ND | 2.776285482 | 0.000187179 | DG | Polyunsaturated | 37 |
| TG 53:1\|TG 17:0_18:0_18:1_[M+NH4]+_25.771 | Control HFD - Control ND | 2.82715892 | 0.000275436 | TG | Monounsaturated | 53 |
| TG 53:1\|TG 17:0_18:0_18:1_[M+NH4]+_25.771 | Control HFD - Control ND | 2.82715892 | 0.000275436 | TG | Monounsaturated | 53 |
| PE 38:5_[M+H]+_16.594 | Control HFD - Control ND | 2.883780772 | 5.86892E-07 | PE | Polyunsaturated | 38 |
| PE 38:5_[M+H]+_16.594 | Control HFD - Control ND | 2.883780772 | 5.86892E-07 | PE | Polyunsaturated | 38 |
| CE 17:0_[M+NH4]+_26.279 | Control HFD - Control ND | 3.113805825 | 0.047699729 | CE | Saturated | 17 |
| CE 17:0_[M+NH4]+_26.279 | Control HFD - Control ND | 3.113805825 | 0.047699729 | CE | Saturated | 17 |
| FAHFA 25:0;O\|FAHFA 16:0/9:0;O_[M-H]-_13.894 | Control HFD - Control ND | 3.193427009 | 0.006487885 | FAHFA | Saturated | 25 |
| FAHFA 25:0;O\|FAHFA 16:0/9:0;O_[M-H]-_13.894 | Control HFD - Control ND | 3.193427009 | 0.006487885 | FAHFA | Saturated | 25 |
| TG 53:2\|TG 16:0_18:1_19:1_[M+NH4]+_25.752 | Control HFD - Control ND | 3.404051264 | 1.87833E-06 | TG | Polyunsaturated | 53 |
| TG 53:2\|TG 16:0_18:1_19:1_[M+NH4]+_25.752 | Control HFD - Control ND | 3.404051264 | 1.87833E-06 | TG | Polyunsaturated | 53 |
| TG 45:0\|TG 14:0_15:0_16:0_[M+NH4]+_23.976 | Control HFD - Control ND | 3.480099887 | 0.017237638 | TG | Saturated | 45 |
| TG 45:0\|TG 14:0_15:0_16:0_[M+NH4]+_23.976 | Control HFD - Control ND | 3.480099887 | 0.017237638 | TG | Saturated | 45 |
| DG 39:6\|DG 17:0_22:6_[M+NH4]+_18.284 | Control HFD - Control ND | 3.485572834 | 0.00275847 | DG | Polyunsaturated | 39 |
| DG 39:6\|DG 17:0_22:6_[M+NH4]+_18.284 | Control HFD - Control ND | 3.485572834 | 0.00275847 | DG | Polyunsaturated | 39 |
| TG 45:1\|TG 14:0_15:0_16:1_[M+NH4]+_23.221 | Control HFD - Control ND | 3.604239497 | 0.000627914 | TG | Monounsaturated | 45 |
| TG 45:1\|TG 14:0_15:0_16:1_[M+NH4]+_23.221 | Control HFD - Control ND | 3.604239497 | 0.000627914 | TG | Monounsaturated | 45 |
| TG 52:4;1O\|TG 17:1_17:1_18:2;1O_[M+NH4]+_23.389 | Control HFD - Control ND | 3.743089269 | 0.031737763 | OxTG | Polyunsaturated | 52 |

| | | | | | | |
|---|---|---|---|---|---|---|
| TG 52:4;1O\|TG 17:1_17:1_18:2;1O_[M+NH4]+_23.389 | Control HFD - Control ND | 3.743089269 | 0.031737763 | OxTG | Polyunsaturated | 52 |
| SM 36:2;2O\|SM 18:2;2O/18:0_[M+H]+_15.791 | Control HFD - Control ND | 3.764914377 | 3.61205E-05 | SM | Polyunsaturated | 36 |
| SM 36:2;2O\|SM 18:2;2O/18:0_[M+H]+_15.791 | Control HFD - Control ND | 3.764914377 | 3.61205E-05 | SM | Polyunsaturated | 36 |
| TG 47:1\|TG 14:0_15:0_18:1_[M+NH4]+_23.931 | Control HFD - Control ND | 3.782583555 | 1.66622E-05 | TG | Monounsaturated | 47 |
| TG 47:1\|TG 14:0_15:0_18:1_[M+NH4]+_23.931 | Control HFD - Control ND | 3.782583555 | 1.66622E-05 | TG | Monounsaturated | 47 |
| DG 35:1\|DG 17:0_18:1_[M+NH4]+_19.727 | Control HFD - Control ND | 4.276284422 | 1.36717E-05 | DG | Monounsaturated | 35 |
| DG 35:1\|DG 17:0_18:1_[M+NH4]+_19.727 | Control HFD - Control ND | 4.276284422 | 1.36717E-05 | DG | Monounsaturated | 35 |
| TG 45:2\|TG 16:0_13:1_16:1_[M+NH4]+_22.953 | Control HFD - Control ND | 4.28507238 | 0.000116475 | TG | Polyunsaturated | 45 |
| TG 45:2\|TG 16:0_13:1_16:1_[M+NH4]+_22.953 | Control HFD - Control ND | 4.28507238 | 0.000116475 | TG | Polyunsaturated | 45 |
| FAHFA 27:0;O\|FAHFA 18:0/9:0;O_[M-H]-_15.809 | Control HFD - Control ND | 4.461784085 | 0.003427595 | FAHFA | Saturated | 27 |
| FAHFA 27:0;O\|FAHFA 18:0/9:0;O_[M-H]-_15.809 | Control HFD - Control ND | 4.461784085 | 0.003427595 | FAHFA | Saturated | 27 |
| TG 51:2\|TG 16:0_17:1_18:1_[M+NH4]+_25.26 | Control HFD - Control ND | 4.794417741 | 2.08449E-09 | TG | Polyunsaturated | 51 |
| TG 51:2\|TG 16:0_17:1_18:1_[M+NH4]+_25.26 | Control HFD - Control ND | 4.794417741 | 2.08449E-09 | TG | Polyunsaturated | 51 |
| TG 71:4;O2\|TG 18:1_18:1_18:1;O(FA 17:0)_[M+NH4]+_26.582 | Control HFD - Control ND | 5.388129173 | 0.013881303 | TG_EST | Polyunsaturated | 71 |
| TG 71:4;O2\|TG 18:1_18:1_18:1;O(FA 17:0)_[M+NH4]+_26.582 | Control HFD - Control ND | 5.388129173 | 0.013881303 | TG_EST | Polyunsaturated | 71 |
| PG 38:1\|PG 20:0_18:1_[M-H]-_16.778 | Control HFD - Control ND | 6.91214449 | 0.009193249 | PG | Monounsaturated | 38 |
| PG 38:1\|PG 20:0_18:1_[M-H]-_16.778 | Control HFD - Control ND | 6.91214449 | 0.009193249 | PG | Monounsaturated | 38 |
| CE 16:1_[M+NH4]+_25.434 | LGR4/5dLKO HFD - LGR4/5dLKO ND | -4.979665441 | 0.001957632 | CE | Monounsaturated | 16 |

| | | | | | | |
|---|---|---|---|---|---|---|
| CE 16:1_[M+NH4]+_25.434 | LGR4/5dLKO HFD - LGR4/5dLKO ND | -4.979665441 | 0.001957632 | CE | Monounsaturated | 16 |
| CE 22:6_[M+NH4]+_24.694 | LGR4/5dLKO HFD - LGR4/5dLKO ND | -4.830297829 | 0.001549422 | CE | Polyunsaturated | 22 |
| CE 22:6_[M+NH4]+_24.694 | LGR4/5dLKO HFD - LGR4/5dLKO ND | -4.830297829 | 0.001549422 | CE | Polyunsaturated | 22 |
| CE 18:2_[M+NH4]+_25.468 | LGR4/5dLKO HFD - LGR4/5dLKO ND | -4.155237569 | 3.86475E-07 | CE | Polyunsaturated | 18 |
| CE 18:2_[M+NH4]+_25.468 | LGR4/5dLKO HFD - LGR4/5dLKO ND | -4.155237569 | 3.86475E-07 | CE | Polyunsaturated | 18 |
| PE 42:6_[M+H]+_18.582 | LGR4/5dLKO HFD - LGR4/5dLKO ND | -4.138068372 | 0.011680666 | PE | Polyunsaturated | 42 |
| PE 42:6_[M+H]+_18.582 | LGR4/5dLKO HFD - LGR4/5dLKO ND | -4.138068372 | 0.011680666 | PE | Polyunsaturated | 42 |
| PE 36:6\|PE 16:1_20:5_[M-H]-_13.782 | LGR4/5dLKO HFD - LGR4/5dLKO ND | -4.039625291 | 5.76123E-10 | PE | Polyunsaturated | 36 |
| PE 36:6\|PE 16:1_20:5_[M-H]-_13.782 | LGR4/5dLKO HFD - LGR4/5dLKO ND | -4.039625291 | 5.76123E-10 | PE | Polyunsaturated | 36 |
| PC 36:6\|PC 16:1_20:5_[M+HCOO]-_13.54 | LGR4/5dLKO HFD - LGR4/5dLKO ND | -4.00312478 | 1.12173E-09 | PC | Polyunsaturated | 36 |
| PC 36:6\|PC 16:1_20:5_[M+HCOO]-_13.54 | LGR4/5dLKO HFD - LGR4/5dLKO ND | -4.00312478 | 1.12173E-09 | PC | Polyunsaturated | 36 |
| PE 36:6_[M+H]+_13.792 | LGR4/5dLKO HFD - LGR4/5dLKO ND | -3.859008799 | 2.91827E-09 | PE | Polyunsaturated | 36 |
| PE 36:6_[M+H]+_13.792 | LGR4/5dLKO HFD - LGR4/5dLKO ND | -3.859008799 | 2.91827E-09 | PE | Polyunsaturated | 36 |
| PE 36:5\|PE 16:0_20:5_[M-H]-_15.295 | LGR4/5dLKO HFD - LGR4/5dLKO ND | -3.502076774 | 1.99265E-11 | PE | Polyunsaturated | 36 |
| PE 36:5\|PE 16:0_20:5_[M-H]-_15.295 | LGR4/5dLKO HFD - LGR4/5dLKO ND | -3.502076774 | 1.99265E-11 | PE | Polyunsaturated | 36 |
| PE 36:5_[M+H]+_15.288 | LGR4/5dLKO HFD - LGR4/5dLKO ND | -3.45663515 | 2.5902E-10 | PE | Polyunsaturated | 36 |
| PE 36:5_[M+H]+_15.288 | LGR4/5dLKO HFD - LGR4/5dLKO ND | -3.45663515 | 2.5902E-10 | PE | Polyunsaturated | 36 |
| CE 24:5_[M+NH4]+_25.693 | LGR4/5dLKO HFD - LGR4/5dLKO ND | -3.174836109 | 0.008664096 | CE | Polyunsaturated | 24 |
| CE 24:5_[M+NH4]+_25.693 | LGR4/5dLKO HFD - LGR4/5dLKO ND | -3.174836109 | 0.008664096 | CE | Polyunsaturated | 24 |
| PI 36:3\|PI 16:0_20:3_[M-H]-_15.261 | LGR4/5dLKO HFD - LGR4/5dLKO ND | -3.126864595 | 1.26149E-06 | PI | Polyunsaturated | 36 |
| PI 36:3\|PI 16:0_20:3_[M-H]-_15.261 | LGR4/5dLKO HFD - LGR4/5dLKO ND | -3.126864595 | 1.26149E-06 | PI | Polyunsaturated | 36 |
| PI 40:6\|PI 18:0_22:6_[M-H]-_15.927 | LGR4/5dLKO HFD - LGR4/5dLKO ND | -3.063746352 | 2.15703E-09 | PI | Polyunsaturated | 40 |
| PI 40:6\|PI 18:0_22:6_[M-H]-_15.927 | LGR4/5dLKO HFD - LGR4/5dLKO ND | -3.063746352 | 2.15703E-09 | PI | Polyunsaturated | 40 |
| PI 32:1;3O\|PI 16:0_16:1;3O_[M-H]-_13.687 | LGR4/5dLKO HFD - LGR4/5dLKO ND | -3.021167451 | 1.61036E-09 | OxPI | Monounsaturated | 32 |

| | | | | | | |
|---|---|---|---|---|---|---|
| PI 32:1;3O\|PI 16:0_16:1;3O_[M-H]-_13.687 | LGR4/5dLKO HFD - LGR4/5dLKO ND | -3.021167451 | 1.61036E-09 | OxPI | Monounsaturated | 32 |
| PC 38:7\|PC 18:2_20:5_[M+HCOO]-_13.937 | LGR4/5dLKO HFD - LGR4/5dLKO ND | -3.012900083 | 1.84696E-08 | PC | Polyunsaturated | 38 |
| PC 38:7\|PC 18:2_20:5_[M+HCOO]-_13.937 | LGR4/5dLKO HFD - LGR4/5dLKO ND | -3.012900083 | 1.84696E-08 | PC | Polyunsaturated | 38 |
| LPC 16:1/0:0_[M+H]+_2.753 | LGR4/5dLKO HFD - LGR4/5dLKO ND | -2.980482718 | 2.59137E-07 | LPC | Polyunsaturated | 16 |
| LPC 16:1/0:0_[M+H]+_2.753 | LGR4/5dLKO HFD - LGR4/5dLKO ND | -2.980482718 | 2.59137E-07 | LPC | Polyunsaturated | 16 |
| LPC 16:1_[M+HCOO]-_2.851 | LGR4/5dLKO HFD - LGR4/5dLKO ND | -2.945674609 | 5.95469E-08 | LPC | Monounsaturated | 16 |
| LPC 16:1_[M+HCOO]-_2.851 | LGR4/5dLKO HFD - LGR4/5dLKO ND | -2.945674609 | 5.95469E-08 | LPC | Monounsaturated | 16 |
| DMPE 32:1\|DMPE 16:0_16:1_[M-H]-_15.804 | LGR4/5dLKO HFD - LGR4/5dLKO ND | -2.900340469 | 1.18059E-08 | DMPE | Monounsaturated | 32 |
| DMPE 32:1\|DMPE 16:0_16:1_[M-H]-_15.804 | LGR4/5dLKO HFD - LGR4/5dLKO ND | -2.900340469 | 1.18059E-08 | DMPE | Monounsaturated | 32 |
| PC 36:5\|PC 16:0_20:5_[M+HCOO]-_15.021 | LGR4/5dLKO HFD - LGR4/5dLKO ND | -2.882909063 | 7.58933E-09 | PC | Polyunsaturated | 36 |
| PC 36:5\|PC 16:0_20:5_[M+HCOO]-_15.021 | LGR4/5dLKO HFD - LGR4/5dLKO ND | -2.882909063 | 7.58933E-09 | PC | Polyunsaturated | 36 |
| PE 32:1\|PE 16:0_16:1_[M-H]-_16.113 | LGR4/5dLKO HFD - LGR4/5dLKO ND | -2.878766263 | 6.25622E-07 | PE | Monounsaturated | 32 |
| PE 32:1\|PE 16:0_16:1_[M-H]-_16.113 | LGR4/5dLKO HFD - LGR4/5dLKO ND | -2.878766263 | 6.25622E-07 | PE | Monounsaturated | 32 |
| CAR 4:0_[M+H]+_1.078 | LGR4/5dLKO HFD - LGR4/5dLKO ND | -2.877043575 | 2.29458E-06 | CAR | Saturated | 4 |
| CAR 4:0_[M+H]+_1.078 | LGR4/5dLKO HFD - LGR4/5dLKO ND | -2.877043575 | 2.29458E-06 | CAR | Saturated | 4 |
| PC 34:3\|PC 16:1_18:2_[M+HCOO]-_14.813 | LGR4/5dLKO HFD - LGR4/5dLKO ND | -2.870233206 | 5.35804E-10 | PC | Polyunsaturated | 34 |
| PC 34:3\|PC 16:1_18:2_[M+HCOO]-_14.813 | LGR4/5dLKO HFD - LGR4/5dLKO ND | -2.870233206 | 5.35804E-10 | PC | Polyunsaturated | 34 |
| DMPE 36:5\|DMPE 16:0_20:5_[M-H]-_15.001 | LGR4/5dLKO HFD - LGR4/5dLKO ND | -2.83860708 | 1.59928E-06 | DMPE | Polyunsaturated | 36 |
| DMPE 36:5\|DMPE 16:0_20:5_[M-H]-_15.001 | LGR4/5dLKO HFD - LGR4/5dLKO ND | -2.83860708 | 1.59928E-06 | DMPE | Polyunsaturated | 36 |
| DMPE 34:3\|DMPE 16:1_18:2_[M-H]-_14.798 | LGR4/5dLKO HFD - LGR4/5dLKO ND | -2.82687799 | 1.51092E-09 | DMPE | Polyunsaturated | 34 |
| DMPE 34:3\|DMPE 16:1_18:2_[M-H]-_14.798 | LGR4/5dLKO HFD - LGR4/5dLKO ND | -2.82687799 | 1.51092E-09 | DMPE | Polyunsaturated | 34 |
| PI 34:2\|PI 16:0_18:2_[M-H]-_14.837 | LGR4/5dLKO HFD - LGR4/5dLKO ND | -2.805872923 | 6.59724E-06 | PI | Polyunsaturated | 34 |
| PI 34:2\|PI 16:0_18:2_[M-H]-_14.837 | LGR4/5dLKO HFD - LGR4/5dLKO ND | -2.805872923 | 6.59724E-06 | PI | Polyunsaturated | 34 |
| PS 36:5;3O\|PS 20:4_16:1;3O_[M-H]-_14.512 | LGR4/5dLKO HFD - LGR4/5dLKO ND | -2.803481133 | 1.33781E-07 | OxPS | Polyunsaturated | 36 |

| | | | | | | |
|---|---|---|---|---|---|---|
| PS 36:5;3O\|PS 20:4_16:1;3O_[M-H]-_14.512 | LGR4/5dLKO HFD - LGR4/5dLKO ND | -2.803481133 | 1.33781E-07 | OxPS | Polyunsaturated | 36 |
| PE 38:7\|PE 16:1_22:6_[M+H]+_14.5 | LGR4/5dLKO HFD - LGR4/5dLKO ND | -2.78462709 | 9.24275E-08 | PE | Polyunsaturated | 38 |
| PE 38:7\|PE 16:1_22:6_[M+H]+_14.5 | LGR4/5dLKO HFD - LGR4/5dLKO ND | -2.78462709 | 9.24275E-08 | PE | Polyunsaturated | 38 |
| PE 38:7\|PE 16:1_22:6_[M-H]-_14.503 | LGR4/5dLKO HFD - LGR4/5dLKO ND | -2.772080527 | 1.24268E-08 | PE | Polyunsaturated | 38 |
| PE 38:7\|PE 16:1_22:6_[M-H]-_14.503 | LGR4/5dLKO HFD - LGR4/5dLKO ND | -2.772080527 | 1.24268E-08 | PE | Polyunsaturated | 38 |
| CAR 6:0_[M+H]+_1.129 | LGR4/5dLKO HFD - LGR4/5dLKO ND | -2.759035494 | 6.69104E-05 | CAR | Saturated | 6 |
| CAR 6:0_[M+H]+_1.129 | LGR4/5dLKO HFD - LGR4/5dLKO ND | -2.759035494 | 6.69104E-05 | CAR | Saturated | 6 |
| PC 32:1\|PC 16:0_16:1_[M+HCOO]-_15.806 | LGR4/5dLKO HFD - LGR4/5dLKO ND | -2.690107403 | 5.50231E-09 | PC | Monounsaturated | 32 |
| PC 32:1\|PC 16:0_16:1_[M+HCOO]-_15.806 | LGR4/5dLKO HFD - LGR4/5dLKO ND | -2.690107403 | 5.50231E-09 | PC | Monounsaturated | 32 |
| PE 38:7\|PE 18:2_20:5_[M-H]-_14.177 | LGR4/5dLKO HFD - LGR4/5dLKO ND | -2.645006858 | 5.88919E-07 | PE | Polyunsaturated | 38 |
| PE 38:7\|PE 18:2_20:5_[M-H]-_14.177 | LGR4/5dLKO HFD - LGR4/5dLKO ND | -2.645006858 | 5.88919E-07 | PE | Polyunsaturated | 38 |
| DMPE 38:7\|DMPE 16:1_22:6_[M-H]-_14.254 | LGR4/5dLKO HFD - LGR4/5dLKO ND | -2.623573151 | 8.8577E-07 | DMPE | Polyunsaturated | 38 |
| DMPE 38:7\|DMPE 16:1_22:6_[M-H]-_14.254 | LGR4/5dLKO HFD - LGR4/5dLKO ND | -2.623573151 | 8.8577E-07 | DMPE | Polyunsaturated | 38 |
| PC 38:7\|PC 16:1_22:6_[M+HCOO]-_14.253 | LGR4/5dLKO HFD - LGR4/5dLKO ND | -2.608507912 | 1.0798E-06 | PC | Polyunsaturated | 38 |
| PC 38:7\|PC 16:1_22:6_[M+HCOO]-_14.253 | LGR4/5dLKO HFD - LGR4/5dLKO ND | -2.608507912 | 1.0798E-06 | PC | Polyunsaturated | 38 |
| PI 40:7\|PI 18:1_22:6_[M-H]-_14.542 | LGR4/5dLKO HFD - LGR4/5dLKO ND | -2.600444472 | 8.497E-08 | PI | Polyunsaturated | 40 |
| PI 40:7\|PI 18:1_22:6_[M-H]-_14.542 | LGR4/5dLKO HFD - LGR4/5dLKO ND | -2.600444472 | 8.497E-08 | PI | Polyunsaturated | 40 |
| PI 38:3_[M+NH4]+_16.73 | LGR4/5dLKO HFD - LGR4/5dLKO ND | -2.599840001 | 4.85757E-10 | PI | Polyunsaturated | 38 |
| PI 38:3_[M+NH4]+_16.73 | LGR4/5dLKO HFD - LGR4/5dLKO ND | -2.599840001 | 4.85757E-10 | PI | Polyunsaturated | 38 |
| PE 36:5;3O\|PE 20:4_16:1;3O_[M-H]-_13.841 | LGR4/5dLKO HFD - LGR4/5dLKO ND | -2.580620963 | 1.89344E-05 | OxPE | Polyunsaturated | 36 |
| PE 36:5;3O\|PE 20:4_16:1;3O_[M-H]-_13.841 | LGR4/5dLKO HFD - LGR4/5dLKO ND | -2.580620963 | 1.89344E-05 | OxPE | Polyunsaturated | 36 |
| LPC 16:1/0:0_[M+H]+_2.811 | LGR4/5dLKO HFD - LGR4/5dLKO ND | -2.563878918 | 1.93925E-06 | LPC | Polyunsaturated | 16 |
| LPC 16:1/0:0_[M+H]+_2.811 | LGR4/5dLKO HFD - LGR4/5dLKO ND | -2.563878918 | 1.93925E-06 | LPC | Polyunsaturated | 16 |
| LPC 20:0_[M+HCOO]-_9.029 | LGR4/5dLKO HFD - LGR4/5dLKO ND | -2.516734805 | 0.000213785 | LPC | Saturated | 20 |

| | | | | | | |
|---|---|---|---|---|---|---|
| LPC 20:0_[M+HCOO]-_9.029 | LGR4/5dLKO HFD - LGR4/5dLKO ND | -2.516734805 | 0.000213785 | LPC | Saturated | 20 |
| LPC 16:1/0:0_[M+H]+_2.85 | LGR4/5dLKO HFD - LGR4/5dLKO ND | -2.506807071 | 3.2199E-06 | LPC | Polyunsaturated | 16 |
| LPC 16:1/0:0_[M+H]+_2.85 | LGR4/5dLKO HFD - LGR4/5dLKO ND | -2.506807071 | 3.2199E-06 | LPC | Polyunsaturated | 16 |
| PE 34:3\|PE 16:1_18:2_[M-H]-_15.052 | LGR4/5dLKO HFD - LGR4/5dLKO ND | -2.479196166 | 5.28057E-06 | PE | Polyunsaturated | 34 |
| PE 34:3\|PE 16:1_18:2_[M-H]-_15.052 | LGR4/5dLKO HFD - LGR4/5dLKO ND | -2.479196166 | 5.28057E-06 | PE | Polyunsaturated | 34 |
| PE 32:2\|PE 16:1_16:1_[M-H]-_14.665 | LGR4/5dLKO HFD - LGR4/5dLKO ND | -2.470959896 | 0.000234304 | PE | Polyunsaturated | 32 |
| PE 32:2\|PE 16:1_16:1_[M-H]-_14.665 | LGR4/5dLKO HFD - LGR4/5dLKO ND | -2.470959896 | 0.000234304 | PE | Polyunsaturated | 32 |
| PE 36:6\|PE 14:0_22:6_[M-H]-_14.259 | LGR4/5dLKO HFD - LGR4/5dLKO ND | -2.403343558 | 1.07025E-07 | PE | Polyunsaturated | 36 |
| PE 36:6\|PE 14:0_22:6_[M-H]-_14.259 | LGR4/5dLKO HFD - LGR4/5dLKO ND | -2.403343558 | 1.07025E-07 | PE | Polyunsaturated | 36 |
| PC 32:2\|PC 16:1_16:1_[M+HCOO]-_14.521 | LGR4/5dLKO HFD - LGR4/5dLKO ND | -2.400609404 | 2.11589E-08 | PC | Polyunsaturated | 32 |
| PC 32:2\|PC 16:1_16:1_[M+HCOO]-_14.521 | LGR4/5dLKO HFD - LGR4/5dLKO ND | -2.400609404 | 2.11589E-08 | PC | Polyunsaturated | 32 |
| PS 38:6\|PS 16:0_22:6_[M-H]-_14.544 | LGR4/5dLKO HFD - LGR4/5dLKO ND | -2.388157112 | 9.1771E-08 | PS | Polyunsaturated | 38 |
| PS 38:6\|PS 16:0_22:6_[M-H]-_14.544 | LGR4/5dLKO HFD - LGR4/5dLKO ND | -2.388157112 | 9.1771E-08 | PS | Polyunsaturated | 38 |
| LPE 16:0_[M-H]-_4.132 | LGR4/5dLKO HFD - LGR4/5dLKO ND | -2.359702773 | 2.32899E-08 | LPE | Saturated | 16 |
| LPE 16:0_[M-H]-_4.132 | LGR4/5dLKO HFD - LGR4/5dLKO ND | -2.359702773 | 2.32899E-08 | LPE | Saturated | 16 |
| DMPE 44:12\|DMPE 22:6_22:6_[M-H]-_16.346 | LGR4/5dLKO HFD - LGR4/5dLKO ND | -2.359494909 | 1.0824E-05 | DMPE | Polyunsaturated | 44 |
| DMPE 44:12\|DMPE 22:6_22:6_[M-H]-_16.346 | LGR4/5dLKO HFD - LGR4/5dLKO ND | -2.359494909 | 1.0824E-05 | DMPE | Polyunsaturated | 44 |
| PC 34:4\|PC 16:1_18:3_[M+HCOO]-_13.785 | LGR4/5dLKO HFD - LGR4/5dLKO ND | -2.358104422 | 0.012927082 | PC | Polyunsaturated | 34 |
| PC 34:4\|PC 16:1_18:3_[M+HCOO]-_13.785 | LGR4/5dLKO HFD - LGR4/5dLKO ND | -2.358104422 | 0.012927082 | PC | Polyunsaturated | 34 |
| PE 36:5\|PE 18:2_18:3_[M-H]-_14.364 | LGR4/5dLKO HFD - LGR4/5dLKO ND | -2.356989809 | 2.2183E-05 | PE | Polyunsaturated | 36 |
| PE 36:5\|PE 18:2_18:3_[M-H]-_14.364 | LGR4/5dLKO HFD - LGR4/5dLKO ND | -2.356989809 | 2.2183E-05 | PE | Polyunsaturated | 36 |
| PE 34:3\|PE 16:1_18:2_[M+H]+_15.04 | LGR4/5dLKO HFD - LGR4/5dLKO ND | -2.335743543 | 1.28641E-05 | PE | Polyunsaturated | 34 |
| PE 34:3\|PE 16:1_18:2_[M+H]+_15.04 | LGR4/5dLKO HFD - LGR4/5dLKO ND | -2.335743543 | 1.28641E-05 | PE | Polyunsaturated | 34 |
| PC 40:9\|PC 20:4_20:5_[M+HCOO]-_13.628 | LGR4/5dLKO HFD - LGR4/5dLKO ND | -2.329541198 | 3.20247E-05 | PC | Polyunsaturated | 40 |

| | | | | | | |
|---|---|---|---|---|---|---|
| PC 40:9\|PC 20:4_20:5_[M+HCOO]-_13.628 | LGR4/5dLKO HFD - LGR4/5dLKO ND | -2.329541198 | 3.20247E-05 | PC | Polyunsaturated | 40 |
| PC 42:6\|PC 20:0_22:6_[M+HCOO]-_18.453 | LGR4/5dLKO HFD - LGR4/5dLKO ND | -2.317814573 | 0.011697284 | PC | Polyunsaturated | 42 |
| PC 42:6\|PC 20:0_22:6_[M+HCOO]-_18.453 | LGR4/5dLKO HFD - LGR4/5dLKO ND | -2.317814573 | 0.011697284 | PC | Polyunsaturated | 42 |
| PS 38:6_[M+H]+_14.544 | LGR4/5dLKO HFD - LGR4/5dLKO ND | -2.30440765 | 6.06874E-07 | PS | Polyunsaturated | 38 |
| PS 38:6_[M+H]+_14.544 | LGR4/5dLKO HFD - LGR4/5dLKO ND | -2.30440765 | 6.06874E-07 | PS | Polyunsaturated | 38 |
| TG 60:13\|TG 16:1_22:6_22:6_[M+NH4]+_22.056 | LGR4/5dLKO HFD - LGR4/5dLKO ND | -2.262379819 | 0.025436882 | TG | Polyunsaturated | 60 |
| TG 60:13\|TG 16:1_22:6_22:6_[M+NH4]+_22.056 | LGR4/5dLKO HFD - LGR4/5dLKO ND | -2.262379819 | 0.025436882 | TG | Polyunsaturated | 60 |
| PC 42:6_[M+H]+_18.414 | LGR4/5dLKO HFD - LGR4/5dLKO ND | -2.260075313 | 0.016127558 | PC | Polyunsaturated | 42 |
| PC 42:6_[M+H]+_18.414 | LGR4/5dLKO HFD - LGR4/5dLKO ND | -2.260075313 | 0.016127558 | PC | Polyunsaturated | 42 |
| PC 30:1\|PC 14:0_16:1_[M+HCOO]-_14.157 | LGR4/5dLKO HFD - LGR4/5dLKO ND | -2.236152694 | 4.48585E-07 | PC | Monounsaturated | 30 |
| PC 30:1\|PC 14:0_16:1_[M+HCOO]-_14.157 | LGR4/5dLKO HFD - LGR4/5dLKO ND | -2.236152694 | 4.48585E-07 | PC | Monounsaturated | 30 |
| PI 38:3\|PI 18:0_20:3_[M-H]-_16.729 | LGR4/5dLKO HFD - LGR4/5dLKO ND | -2.235677971 | 6.78904E-09 | PI | Polyunsaturated | 38 |
| PI 38:3\|PI 18:0_20:3_[M-H]-_16.729 | LGR4/5dLKO HFD - LGR4/5dLKO ND | -2.235677971 | 6.78904E-09 | PI | Polyunsaturated | 38 |
| CL 78:10\|CL 18:1_18:1_22:3_20:5_[M-2H]2-_15.417 | LGR4/5dLKO HFD - LGR4/5dLKO ND | -2.228217812 | 4.67862E-09 | CL | Polyunsaturated | 78 |
| CL 78:10\|CL 18:1_18:1_22:3_20:5_[M-2H]2-_15.417 | LGR4/5dLKO HFD - LGR4/5dLKO ND | -2.228217812 | 4.67862E-09 | CL | Polyunsaturated | 78 |
| PE 38:6\|PE 18:1_20:5_[M-H]-_15.42 | LGR4/5dLKO HFD - LGR4/5dLKO ND | -2.224183638 | 6.94657E-09 | PE | Polyunsaturated | 38 |
| PE 38:6\|PE 18:1_20:5_[M-H]-_15.42 | LGR4/5dLKO HFD - LGR4/5dLKO ND | -2.224183638 | 6.94657E-09 | PE | Polyunsaturated | 38 |
| LPC 16:0/0:0_[M+H]+_4.017 | LGR4/5dLKO HFD - LGR4/5dLKO ND | -2.196429562 | 5.17388E-09 | LPC | NA | 16 |
| LPC 16:0/0:0_[M+H]+_4.017 | LGR4/5dLKO HFD - LGR4/5dLKO ND | -2.196429562 | 5.17388E-09 | LPC | NA | 16 |
| PI 36:3\|PI 18:1_18:2_[M-H]-_15.003 | LGR4/5dLKO HFD - LGR4/5dLKO ND | -2.193107618 | 8.54448E-05 | PI | Polyunsaturated | 36 |
| PI 36:3\|PI 18:1_18:2_[M-H]-_15.003 | LGR4/5dLKO HFD - LGR4/5dLKO ND | -2.193107618 | 8.54448E-05 | PI | Polyunsaturated | 36 |
| LPE 16:0_[M+H]+_4.113 | LGR4/5dLKO HFD - LGR4/5dLKO ND | -2.186012868 | 5.15738E-08 | LPE | Saturated | 16 |
| LPE 16:0_[M+H]+_4.113 | LGR4/5dLKO HFD - LGR4/5dLKO ND | -2.186012868 | 5.15738E-08 | LPE | Saturated | 16 |
| LPE 16:1_[M-H]-_2.902 | LGR4/5dLKO HFD - LGR4/5dLKO ND | -2.179640862 | 1.53036E-06 | LPE | Monounsaturated | 16 |

| | | | | | | |
|---|---|---|---|---|---|---|
| LPE 16:1_[M-H]-_2.902 | LGR4/5dLKO HFD - LGR4/5dLKO ND | -2.179640862 | 1.53036E-06 | LPE | Monounsaturated | 16 |
| PC 36:5\|PC 18:2_18:3_[M+HCOO]-_14.146 | LGR4/5dLKO HFD - LGR4/5dLKO ND | -2.157887655 | 0.004190198 | PC | Polyunsaturated | 36 |
| PC 36:5\|PC 18:2_18:3_[M+HCOO]-_14.146 | LGR4/5dLKO HFD - LGR4/5dLKO ND | -2.157887655 | 0.004190198 | PC | Polyunsaturated | 36 |
| PE 34:2\|PE 16:0_18:2_[M+H]+_16.433 | LGR4/5dLKO HFD - LGR4/5dLKO ND | -2.145235188 | 3.94475E-09 | PE | Polyunsaturated | 34 |
| PE 34:2\|PE 16:0_18:2_[M+H]+_16.433 | LGR4/5dLKO HFD - LGR4/5dLKO ND | -2.145235188 | 3.94475E-09 | PE | Polyunsaturated | 34 |
| PE O-37:2\|PE O-19:0_18:2_[M-H]-_16.14 | LGR4/5dLKO HFD - LGR4/5dLKO ND | -2.132140889 | 9.08646E-07 | EtherPE | Polyunsaturated | |
| PE O-37:2\|PE O-19:0_18:2_[M-H]-_16.14 | LGR4/5dLKO HFD - LGR4/5dLKO ND | -2.132140889 | 9.08646E-07 | EtherPE | Polyunsaturated | |
| LPC 18:1/0:0_[M+H]+_4.372 | LGR4/5dLKO HFD - LGR4/5dLKO ND | -2.120142441 | 3.33401E-07 | LPC | Polyunsaturated | 18 |
| LPC 18:1/0:0_[M+H]+_4.372 | LGR4/5dLKO HFD - LGR4/5dLKO ND | -2.120142441 | 3.33401E-07 | LPC | Polyunsaturated | 18 |
| LPC 16:0_[M+HCOO]-_4.059 | LGR4/5dLKO HFD - LGR4/5dLKO ND | -2.101206493 | 1.37543E-09 | LPC | Saturated | 16 |
| LPC 16:0_[M+HCOO]-_4.059 | LGR4/5dLKO HFD - LGR4/5dLKO ND | -2.101206493 | 1.37543E-09 | LPC | Saturated | 16 |
| PE 36:5\|PE 16:1_20:4_[M-H]-_14.849 | LGR4/5dLKO HFD - LGR4/5dLKO ND | -2.100674265 | 4.2079E-07 | PE | Polyunsaturated | 36 |
| PE 36:5\|PE 16:1_20:4_[M-H]-_14.849 | LGR4/5dLKO HFD - LGR4/5dLKO ND | -2.100674265 | 4.2079E-07 | PE | Polyunsaturated | 36 |
| PC 44:12\|PC 22:6_22:6_[M+HCOO]-_13.985 | LGR4/5dLKO HFD - LGR4/5dLKO ND | -2.095788482 | 5.2482E-05 | PC | Polyunsaturated | 44 |
| PC 44:12\|PC 22:6_22:6_[M+HCOO]-_13.985 | LGR4/5dLKO HFD - LGR4/5dLKO ND | -2.095788482 | 5.2482E-05 | PC | Polyunsaturated | 44 |
| PC 36:4_[M+H]+_15.111 | LGR4/5dLKO HFD - LGR4/5dLKO ND | -2.091227424 | 7.55669E-08 | PC | Polyunsaturated | 36 |
| PC 36:4_[M+H]+_15.111 | LGR4/5dLKO HFD - LGR4/5dLKO ND | -2.091227424 | 7.55669E-08 | PC | Polyunsaturated | 36 |
| DG 36:4\|DG 18:1_18:3_[M+NH4]+_17.768 | LGR4/5dLKO HFD - LGR4/5dLKO ND | 2.037913699 | 2.3307E-07 | DG | Polyunsaturated | 36 |
| TG 57:5\|TG 18:1_18:1_21:3_[M+NH4]+_24.868 | LGR4/5dLKO HFD - LGR4/5dLKO ND | 2.0615006 | 3.74427E-08 | TG | Polyunsaturated | 57 |
| DG 40:6\|DG 18:1_22:5_[M+NH4]+_18.362 | LGR4/5dLKO HFD - LGR4/5dLKO ND | 2.079407248 | 3.14235E-06 | DG | Polyunsaturated | 40 |
| TG 58:3\|TG 18:1_20:1_20:1_[M+NH4]+_25.898 | LGR4/5dLKO HFD - LGR4/5dLKO ND | 2.109823759 | 1.07424E-05 | TG | Polyunsaturated | 58 |
| TG 70:4;O2\|TG 16:0_18:1_18:1;O(FA 18:1)_[M+NH4]+_26.402 | LGR4/5dLKO HFD - LGR4/5dLKO ND | 2.11809588 | 0.004803043 | TG_EST | Polyunsaturated | 70 |
| TG 49:5\|TG 15:1_16:1_18:3_[M+NH4]+_22.269 | LGR4/5dLKO HFD - LGR4/5dLKO ND | 2.118712385 | 2.72711E-05 | TG | Polyunsaturated | 49 |
| PE 38:5_[M+H]+_16.594 | LGR4/5dLKO HFD - LGR4/5dLKO ND | 2.159783583 | 7.90823E-08 | PE | Polyunsaturated | 38 |

| | | | | | | |
|---|---|---|---|---|---|---|
| TG 50:6\|TG 16:1_18:2_16:3_[M+NH4]+_22.532 | LGR4/5dLKO HFD - LGR4/5dLKO ND | 2.185083025 | 5.04346E-06 | TG | Polyunsaturated | 50 |
| TG 50:6\|TG 16:1_18:2_16:3_[M+NH4]+_22.532 | LGR4/5dLKO HFD - LGR4/5dLKO ND | 2.185083025 | 5.04346E-06 | TG | Polyunsaturated | 50 |
| TG 61:4\|TG 18:1_25:1_18:2_[M+NH4]+_26.159 | LGR4/5dLKO HFD - LGR4/5dLKO ND | 2.187666738 | 0.002052559 | TG | Polyunsaturated | 61 |
| TG 61:4\|TG 18:1_25:1_18:2_[M+NH4]+_26.159 | LGR4/5dLKO HFD - LGR4/5dLKO ND | 2.187666738 | 0.002052559 | TG | Polyunsaturated | 61 |
| TG 55:6\|TG 16:0_17:0_22:6_[M+NH4]+_24.184 | LGR4/5dLKO HFD - LGR4/5dLKO ND | 2.188330347 | 4.56022E-06 | TG | Polyunsaturated | 55 |
| TG 55:6\|TG 16:0_17:0_22:6_[M+NH4]+_24.184 | LGR4/5dLKO HFD - LGR4/5dLKO ND | 2.188330347 | 4.56022E-06 | TG | Polyunsaturated | 55 |
| TG 51:1\|TG 16:0_17:0_18:1_[M+NH4]+_25.313 | LGR4/5dLKO HFD - LGR4/5dLKO ND | 2.207059926 | 5.21218E-05 | TG | Monounsaturated | 51 |
| TG 51:1\|TG 16:0_17:0_18:1_[M+NH4]+_25.313 | LGR4/5dLKO HFD - LGR4/5dLKO ND | 2.207059926 | 5.21218E-05 | TG | Monounsaturated | 51 |
| TG 54:6;1O\|TG 18:1_18:1_18:4;1O_[M+NH4]+_23.588 | LGR4/5dLKO HFD - LGR4/5dLKO ND | 2.237554202 | 0.000430642 | OxTG | Polyunsaturated | 54 |
| TG 54:6;1O\|TG 18:1_18:1_18:4;1O_[M+NH4]+_23.588 | LGR4/5dLKO HFD - LGR4/5dLKO ND | 2.237554202 | 0.000430642 | OxTG | Polyunsaturated | 54 |
| TG 56:5;1O\|TG 19:0_20:3_17:2;1O_[M+NH4]+_25.33 | LGR4/5dLKO HFD - LGR4/5dLKO ND | 2.260061493 | 6.05112E-10 | OxTG | Polyunsaturated | 56 |
| TG 56:5;1O\|TG 19:0_20:3_17:2;1O_[M+NH4]+_25.33 | LGR4/5dLKO HFD - LGR4/5dLKO ND | 2.260061493 | 6.05112E-10 | OxTG | Polyunsaturated | 56 |
| TG 45:2\|TG 11:0_16:0_18:2_[M+NH4]+_22.638 | LGR4/5dLKO HFD - LGR4/5dLKO ND | 2.282587046 | 0.000920208 | TG | Polyunsaturated | 45 |
| TG 45:2\|TG 11:0_16:0_18:2_[M+NH4]+_22.638 | LGR4/5dLKO HFD - LGR4/5dLKO ND | 2.282587046 | 0.000920208 | TG | Polyunsaturated | 45 |
| TG 49:4\|TG 15:1_16:1_18:2_[M+NH4]+_22.816 | LGR4/5dLKO HFD - LGR4/5dLKO ND | 2.302807514 | 6.3788E-05 | TG | Polyunsaturated | 49 |
| TG 49:4\|TG 15:1_16:1_18:2_[M+NH4]+_22.816 | LGR4/5dLKO HFD - LGR4/5dLKO ND | 2.302807514 | 6.3788E-05 | TG | Polyunsaturated | 49 |
| TG 47:1\|TG 14:0_15:0_18:1_[M+NH4]+_23.931 | LGR4/5dLKO HFD - LGR4/5dLKO ND | 2.304491786 | 7.40414E-05 | TG | Monounsaturated | 47 |
| TG 47:1\|TG 14:0_15:0_18:1_[M+NH4]+_23.931 | LGR4/5dLKO HFD - LGR4/5dLKO ND | 2.304491786 | 7.40414E-05 | TG | Monounsaturated | 47 |
| TG 57:4\|TG 18:1_21:1_18:2_[M+NH4]+_25.33 | LGR4/5dLKO HFD - LGR4/5dLKO ND | 2.321755157 | 5.86516E-10 | TG | Polyunsaturated | 57 |
| TG 57:4\|TG 18:1_21:1_18:2_[M+NH4]+_25.33 | LGR4/5dLKO HFD - LGR4/5dLKO ND | 2.321755157 | 5.86516E-10 | TG | Polyunsaturated | 57 |
| TG 46:3\|TG 14:0_14:1_18:2_[M+NH4]+_22.498 | LGR4/5dLKO HFD - LGR4/5dLKO ND | 2.340399979 | 0.00113733 | TG | Polyunsaturated | 46 |
| TG 46:3\|TG 14:0_14:1_18:2_[M+NH4]+_22.498 | LGR4/5dLKO HFD - LGR4/5dLKO ND | 2.340399979 | 0.00113733 | TG | Polyunsaturated | 46 |
| PE O-37:5\|PE O-17:1_20:4_[M-H]-_17.301 | LGR4/5dLKO HFD - LGR4/5dLKO ND | 2.358835439 | 7.06836E-09 | EtherPE | Polyunsaturated | |
| PE O-37:5\|PE O-17:1_20:4_[M-H]-_17.301 | LGR4/5dLKO HFD - LGR4/5dLKO ND | 2.358835439 | 7.06836E-09 | EtherPE | Polyunsaturated | |

| | | | | | | |
|---|---|---|---|---|---|---|
| DG 40:5\|DG 18:1_22:4_[M+NH4]+_19.103 | LGR4/5dLKO HFD - LGR4/5dLKO ND | 2.423052573 | 1.01041E-07 | DG | Polyunsaturated | 40 |
| DG 40:5\|DG 18:1_22:4_[M+NH4]+_19.103 | LGR4/5dLKO HFD - LGR4/5dLKO ND | 2.423052573 | 1.01041E-07 | DG | Polyunsaturated | 40 |
| TG 52:5;1O\|TG 17:1_17:1_18:3;1O_[M+NH4]+_22.732 | LGR4/5dLKO HFD - LGR4/5dLKO ND | 2.425895777 | 0.001418813 | OxTG | Polyunsaturated | 52 |
| TG 52:5;1O\|TG 17:1_17:1_18:3;1O_[M+NH4]+_22.732 | LGR4/5dLKO HFD - LGR4/5dLKO ND | 2.425895777 | 0.001418813 | OxTG | Polyunsaturated | 52 |
| TG 57:7\|TG 17:0_18:1_22:6_[M+NH4]+_24.198 | LGR4/5dLKO HFD - LGR4/5dLKO ND | 2.443883889 | 1.35592E-06 | TG | Polyunsaturated | 57 |
| TG 57:7\|TG 17:0_18:1_22:6_[M+NH4]+_24.198 | LGR4/5dLKO HFD - LGR4/5dLKO ND | 2.443883889 | 1.35592E-06 | TG | Polyunsaturated | 57 |
| PG 42:6\|PG 20:0_22:6_[M-H]-_15.394 | LGR4/5dLKO HFD - LGR4/5dLKO ND | 2.452706213 | 0.000127712 | PG | Polyunsaturated | 42 |
| PG 42:6\|PG 20:0_22:6_[M-H]-_15.394 | LGR4/5dLKO HFD - LGR4/5dLKO ND | 2.452706213 | 0.000127712 | PG | Polyunsaturated | 42 |
| TG 54:6;1O\|TG 18:1_18:1_18:4;1O_[M+NH4]+_22.77 | LGR4/5dLKO HFD - LGR4/5dLKO ND | 2.539313244 | 0.003965969 | OxTG | Polyunsaturated | 54 |
| TG 54:6;1O\|TG 18:1_18:1_18:4;1O_[M+NH4]+_22.77 | LGR4/5dLKO HFD - LGR4/5dLKO ND | 2.539313244 | 0.003965969 | OxTG | Polyunsaturated | 54 |
| TG 72:4;O2\|TG 18:1_18:1_18:0;O(FA 18:1)_[M+NH4]+_26.733 | LGR4/5dLKO HFD - LGR4/5dLKO ND | 2.553828839 | 0.00517545 | TG_EST | Polyunsaturated | 72 |
| TG 72:4;O2\|TG 18:1_18:1_18:0;O(FA 18:1)_[M+NH4]+_26.733 | LGR4/5dLKO HFD - LGR4/5dLKO ND | 2.553828839 | 0.00517545 | TG_EST | Polyunsaturated | 72 |
| TG 59:2\|TG 18:0_18:1_23:1_[M+NH4]+_26.54 | LGR4/5dLKO HFD - LGR4/5dLKO ND | 2.554012545 | 0.002817108 | TG | Polyunsaturated | 59 |
| TG 59:2\|TG 18:0_18:1_23:1_[M+NH4]+_26.54 | LGR4/5dLKO HFD - LGR4/5dLKO ND | 2.554012545 | 0.002817108 | TG | Polyunsaturated | 59 |
| TG 72:4;O2\|TG 18:1_18:1_18:0;O(FA 18:1)_[M+NH4]+_26.655 | LGR4/5dLKO HFD - LGR4/5dLKO ND | 2.561946268 | 0.008918746 | TG_EST | Polyunsaturated | 72 |
| TG 72:4;O2\|TG 18:1_18:1_18:0;O(FA 18:1)_[M+NH4]+_26.655 | LGR4/5dLKO HFD - LGR4/5dLKO ND | 2.561946268 | 0.008918746 | TG_EST | Polyunsaturated | 72 |
| TG 59:2\|TG 21:0_18:1_20:1_[M+NH4]+_26.502 | LGR4/5dLKO HFD - LGR4/5dLKO ND | 2.568111986 | 0.004868498 | TG | Polyunsaturated | 59 |
| TG 59:2\|TG 21:0_18:1_20:1_[M+NH4]+_26.502 | LGR4/5dLKO HFD - LGR4/5dLKO ND | 2.568111986 | 0.004868498 | TG | Polyunsaturated | 59 |
| DG 36:1\|DG 18:0_18:1_[M+NH4]+_20.457 | LGR4/5dLKO HFD - LGR4/5dLKO ND | 2.569010078 | 9.25868E-05 | DG | Monounsaturated | 36 |
| DG 36:1\|DG 18:0_18:1_[M+NH4]+_20.457 | LGR4/5dLKO HFD - LGR4/5dLKO ND | 2.569010078 | 9.25868E-05 | DG | Monounsaturated | 36 |
| PG 39:4\|PG 18:1_21:3_[M-H]-_15.644 | LGR4/5dLKO HFD - LGR4/5dLKO ND | 2.575828492 | 0.000299099 | PG | Polyunsaturated | 39 |
| PG 39:4\|PG 18:1_21:3_[M-H]-_15.644 | LGR4/5dLKO HFD - LGR4/5dLKO ND | 2.575828492 | 0.000299099 | PG | Polyunsaturated | 39 |
| TG 59:4\|TG 18:1_23:1_18:2_[M+NH4]+_25.787 | LGR4/5dLKO HFD - LGR4/5dLKO ND | 2.606457716 | 5.10356E-06 | TG | Polyunsaturated | 59 |
| TG 59:4\|TG 18:1_23:1_18:2_[M+NH4]+_25.787 | LGR4/5dLKO HFD - LGR4/5dLKO ND | 2.606457716 | 5.10356E-06 | TG | Polyunsaturated | 59 |

| | | | | | | |
|---|---|---|---|---|---|---|
| TG 47:3\|TG 14:1_16:1_17:1_[M+NH4]+_22.706 | LGR4/5dLKO HFD - LGR4/5dLKO ND | 2.62593783 | 7.74489E-05 | TG | Polyunsaturated | 47 |
| TG 47:3\|TG 14:1_16:1_17:1_[M+NH4]+_22.706 | LGR4/5dLKO HFD - LGR4/5dLKO ND | 2.62593783 | 7.74489E-05 | TG | Polyunsaturated | 47 |
| SM 36:2;2O\|SM 18:2;2O/18:0_[M+H]+_15.791 | LGR4/5dLKO HFD - LGR4/5dLKO ND | 2.6433777 | 2.9565E-05 | SM | Polyunsaturated | 36 |
| SM 36:2;2O\|SM 18:2;2O/18:0_[M+H]+_15.791 | LGR4/5dLKO HFD - LGR4/5dLKO ND | 2.6433777 | 2.9565E-05 | SM | Polyunsaturated | 36 |
| TG 45:1\|TG 14:0_15:0_16:1_[M+NH4]+_23.221 | LGR4/5dLKO HFD - LGR4/5dLKO ND | 2.645759614 | 0.000418315 | TG | Monounsaturated | 45 |
| TG 45:1\|TG 14:0_15:0_16:1_[M+NH4]+_23.221 | LGR4/5dLKO HFD - LGR4/5dLKO ND | 2.645759614 | 0.000418315 | TG | Monounsaturated | 45 |
| DG 36:2\|DG 18:1_18:1_[M+NH4]+_19.43 | LGR4/5dLKO HFD - LGR4/5dLKO ND | 2.714685567 | 1.14818E-05 | DG | Polyunsaturated | 36 |
| DG 36:2\|DG 18:1_18:1_[M+NH4]+_19.43 | LGR4/5dLKO HFD - LGR4/5dLKO ND | 2.714685567 | 1.14818E-05 | DG | Polyunsaturated | 36 |
| TG 49:4\|TG 13:1_18:1_18:2_[M+NH4]+_22.956 | LGR4/5dLKO HFD - LGR4/5dLKO ND | 2.72256392 | 5.90529E-08 | TG | Polyunsaturated | 49 |
| TG 49:4\|TG 13:1_18:1_18:2_[M+NH4]+_22.956 | LGR4/5dLKO HFD - LGR4/5dLKO ND | 2.72256392 | 5.90529E-08 | TG | Polyunsaturated | 49 |
| DG 33:1\|DG 15:0_18:1_[M+NH4]+_18.816 | LGR4/5dLKO HFD - LGR4/5dLKO ND | 2.769124461 | 1.43364E-08 | DG | Monounsaturated | 33 |
| DG 33:1\|DG 15:0_18:1_[M+NH4]+_18.816 | LGR4/5dLKO HFD - LGR4/5dLKO ND | 2.769124461 | 1.43364E-08 | DG | Monounsaturated | 33 |
| TG 55:1\|TG 16:0_21:0_18:1_[M+NH4]+_26.172 | LGR4/5dLKO HFD - LGR4/5dLKO ND | 2.793546027 | 0.000961771 | TG | Monounsaturated | 55 |
| TG 55:1\|TG 16:0_21:0_18:1_[M+NH4]+_26.172 | LGR4/5dLKO HFD - LGR4/5dLKO ND | 2.793546027 | 0.000961771 | TG | Monounsaturated | 55 |
| TG 46:4\|TG 14:1_16:1_16:2_[M+NH4]+_21.934 | LGR4/5dLKO HFD - LGR4/5dLKO ND | 2.800494636 | 0.000158348 | TG | Polyunsaturated | 46 |
| TG 46:4\|TG 14:1_16:1_16:2_[M+NH4]+_21.934 | LGR4/5dLKO HFD - LGR4/5dLKO ND | 2.800494636 | 0.000158348 | TG | Polyunsaturated | 46 |
| DG 38:3\|DG 18:1_20:2_[M+NH4]+_19.648 | LGR4/5dLKO HFD - LGR4/5dLKO ND | 2.804903208 | 1.8493E-08 | DG | Polyunsaturated | 38 |
| DG 38:3\|DG 18:1_20:2_[M+NH4]+_19.648 | LGR4/5dLKO HFD - LGR4/5dLKO ND | 2.804903208 | 1.8493E-08 | DG | Polyunsaturated | 38 |
| TG 50:7\|TG 14:0_14:1_22:6_[M+NH4]+_21.934 | LGR4/5dLKO HFD - LGR4/5dLKO ND | 2.80858179 | 0.000277857 | TG | Polyunsaturated | 50 |
| TG 50:7\|TG 14:0_14:1_22:6_[M+NH4]+_21.934 | LGR4/5dLKO HFD - LGR4/5dLKO ND | 2.80858179 | 0.000277857 | TG | Polyunsaturated | 50 |
| TG 59:3\|TG 18:1_18:1_23:1_[M+NH4]+_26.147 | LGR4/5dLKO HFD - LGR4/5dLKO ND | 2.817220393 | 0.000381598 | TG | Polyunsaturated | 59 |
| TG 59:3\|TG 18:1_18:1_23:1_[M+NH4]+_26.147 | LGR4/5dLKO HFD - LGR4/5dLKO ND | 2.817220393 | 0.000381598 | TG | Polyunsaturated | 59 |
| TG 61:3\|TG 18:1_18:1_25:1_[M+NH4]+_26.521 | LGR4/5dLKO HFD - LGR4/5dLKO ND | 2.864297027 | 0.000346293 | TG | Polyunsaturated | 61 |
| TG 61:3\|TG 18:1_18:1_25:1_[M+NH4]+_26.521 | LGR4/5dLKO HFD - LGR4/5dLKO ND | 2.864297027 | 0.000346293 | TG | Polyunsaturated | 61 |

| | | | | | | |
|---|---|---|---|---|---|---|
| TG 57:3\|TG 18:1_19:1_20:1_[M+NH4]+_25.723 | LGR4/5dLKO HFD - LGR4/5dLKO ND | 2.88837171 | 1.79892E-08 | TG | Polyunsaturated | 57 |
| TG 57:3\|TG 18:1_19:1_20:1_[M+NH4]+_25.723 | LGR4/5dLKO HFD - LGR4/5dLKO ND | 2.88837171 | 1.79892E-08 | TG | Polyunsaturated | 57 |
| TG 55:2\|TG 16:0_18:1_21:1_[M+NH4]+_25.735 | LGR4/5dLKO HFD - LGR4/5dLKO ND | 2.889745123 | 3.7848E-08 | TG | Polyunsaturated | 55 |
| TG 55:2\|TG 16:0_18:1_21:1_[M+NH4]+_25.735 | LGR4/5dLKO HFD - LGR4/5dLKO ND | 2.889745123 | 3.7848E-08 | TG | Polyunsaturated | 55 |
| TG 69:4;O2\|TG 16:0_18:1_18:1;O(FA 17:1)_[M+NH4]+_26.251 | LGR4/5dLKO HFD - LGR4/5dLKO ND | 2.901920863 | 0.004385583 | TG_EST | Polyunsaturated | 69 |
| TG 69:4;O2\|TG 16:0_18:1_18:1;O(FA 17:1)_[M+NH4]+_26.251 | LGR4/5dLKO HFD - LGR4/5dLKO ND | 2.901920863 | 0.004385583 | TG_EST | Polyunsaturated | 69 |
| TG 57:3\|TG 21:0_18:1_18:2_[M+NH4]+_25.792 | LGR4/5dLKO HFD - LGR4/5dLKO ND | 2.911998044 | 1.77013E-07 | TG | Polyunsaturated | 57 |
| TG 57:3\|TG 21:0_18:1_18:2_[M+NH4]+_25.792 | LGR4/5dLKO HFD - LGR4/5dLKO ND | 2.911998044 | 1.77013E-07 | TG | Polyunsaturated | 57 |
| TG 45:3\|TG 13:1_14:1_18:1_[M+NH4]+_22.047 | LGR4/5dLKO HFD - LGR4/5dLKO ND | 2.980453116 | 0.003044426 | TG | Polyunsaturated | 45 |
| TG 45:3\|TG 13:1_14:1_18:1_[M+NH4]+_22.047 | LGR4/5dLKO HFD - LGR4/5dLKO ND | 2.980453116 | 0.003044426 | TG | Polyunsaturated | 45 |
| TG 55:6\|TG 16:0_17:1_22:5_[M+NH4]+_24.237 | LGR4/5dLKO HFD - LGR4/5dLKO ND | 2.981694742 | 5.48652E-08 | TG | Polyunsaturated | 55 |
| TG 55:6\|TG 16:0_17:1_22:5_[M+NH4]+_24.237 | LGR4/5dLKO HFD - LGR4/5dLKO ND | 2.981694742 | 5.48652E-08 | TG | Polyunsaturated | 55 |
| TG 53:1\|TG 17:0_18:0_18:1_[M+NH4]+_25.771 | LGR4/5dLKO HFD - LGR4/5dLKO ND | 3.010117996 | 1.66587E-06 | TG | Monounsaturated | 53 |
| TG 53:1\|TG 17:0_18:0_18:1_[M+NH4]+_25.771 | LGR4/5dLKO HFD - LGR4/5dLKO ND | 3.010117996 | 1.66587E-06 | TG | Monounsaturated | 53 |
| TG 72:5;O2\|TG 18:1_18:1_18:0;O(FA 18:2)_[M+NH4]+_26.359 | LGR4/5dLKO HFD - LGR4/5dLKO ND | 3.018823447 | 0.000303678 | TG_EST | Polyunsaturated | 72 |
| TG 72:5;O2\|TG 18:1_18:1_18:0;O(FA 18:2)_[M+NH4]+_26.359 | LGR4/5dLKO HFD - LGR4/5dLKO ND | 3.018823447 | 0.000303678 | TG_EST | Polyunsaturated | 72 |
| TG 50:4;1O\|TG 17:1_17:1_16:2;1O_[M+NH4]+_22.447 | LGR4/5dLKO HFD - LGR4/5dLKO ND | 3.040515183 | 0.000242398 | OxTG | Polyunsaturated | 50 |
| TG 50:4;1O\|TG 17:1_17:1_16:2;1O_[M+NH4]+_22.447 | LGR4/5dLKO HFD - LGR4/5dLKO ND | 3.040515183 | 0.000242398 | OxTG | Polyunsaturated | 50 |
| TG 57:2\|TG 16:0_18:1_23:1_[M+NH4]+_26.142 | LGR4/5dLKO HFD - LGR4/5dLKO ND | 3.040565541 | 0.000634744 | TG | Polyunsaturated | 57 |
| TG 57:2\|TG 16:0_18:1_23:1_[M+NH4]+_26.142 | LGR4/5dLKO HFD - LGR4/5dLKO ND | 3.040565541 | 0.000634744 | TG | Polyunsaturated | 57 |
| TG 59:5\|TG 18:1_19:1_22:3_[M+NH4]+_25.377 | LGR4/5dLKO HFD - LGR4/5dLKO ND | 3.081050091 | 1.94496E-08 | TG | Polyunsaturated | 59 |
| TG 59:5\|TG 18:1_19:1_22:3_[M+NH4]+_25.377 | LGR4/5dLKO HFD - LGR4/5dLKO ND | 3.081050091 | 1.94496E-08 | TG | Polyunsaturated | 59 |
| TG 72:6;O2\|TG 18:1_18:2_18:2;O(FA 18:0)_[M+NH4]+_26.107 | LGR4/5dLKO HFD - LGR4/5dLKO ND | 3.108730737 | 0.005954908 | TG_EST | Polyunsaturated | 72 |
| TG 72:6;O2\|TG 18:1_18:2_18:2;O(FA 18:0)_[M+NH4]+_26.107 | LGR4/5dLKO HFD - LGR4/5dLKO ND | 3.108730737 | 0.005954908 | TG_EST | Polyunsaturated | 72 |

| | | | | | | |
|---|---|---|---|---|---|---|
| DG 38:2\|DG 18:1_20:1_[M+NH4]+_20.42 | LGR4/5dLKO HFD - LGR4/5dLKO ND | 3.185759029 | 2.19477E-08 | DG | Polyunsaturated | 38 |
| DG 38:2\|DG 18:1_20:1_[M+NH4]+_20.42 | LGR4/5dLKO HFD - LGR4/5dLKO ND | 3.185759029 | 2.19477E-08 | DG | Polyunsaturated | 38 |
| DG 42:7\|DG 18:1_24:6_[M+NH4]+_18.686 | LGR4/5dLKO HFD - LGR4/5dLKO ND | 3.206240994 | 9.16334E-08 | DG | Polyunsaturated | 42 |
| DG 42:7\|DG 18:1_24:6_[M+NH4]+_18.686 | LGR4/5dLKO HFD - LGR4/5dLKO ND | 3.206240994 | 9.16334E-08 | DG | Polyunsaturated | 42 |
| DG 35:2\|DG 17:1_18:1_[M+NH4]+_18.874 | LGR4/5dLKO HFD - LGR4/5dLKO ND | 3.262175322 | 2.24968E-09 | DG | Polyunsaturated | 35 |
| DG 35:2\|DG 17:1_18:1_[M+NH4]+_18.874 | LGR4/5dLKO HFD - LGR4/5dLKO ND | 3.262175322 | 2.24968E-09 | DG | Polyunsaturated | 35 |
| TG 51:2;1O\|TG 16:1_18:1_17:0;1O_[M+NH4]+_22.016 | LGR4/5dLKO HFD - LGR4/5dLKO ND | 3.296841792 | 0.016481312 | OxTG | Polyunsaturated | 51 |
| TG 51:2;1O\|TG 16:1_18:1_17:0;1O_[M+NH4]+_22.016 | LGR4/5dLKO HFD - LGR4/5dLKO ND | 3.296841792 | 0.016481312 | OxTG | Polyunsaturated | 51 |
| TG 47:2\|TG 16:0_13:1_18:1_[M+NH4]+_23.634 | LGR4/5dLKO HFD - LGR4/5dLKO ND | 3.347068483 | 7.5515E-09 | TG | Polyunsaturated | 47 |
| TG 47:2\|TG 16:0_13:1_18:1_[M+NH4]+_23.634 | LGR4/5dLKO HFD - LGR4/5dLKO ND | 3.347068483 | 7.5515E-09 | TG | Polyunsaturated | 47 |
| TG 47:4\|TG 13:1_16:1_18:2_[M+NH4]+_22.104 | LGR4/5dLKO HFD - LGR4/5dLKO ND | 3.370552796 | 1.04892E-05 | TG | Polyunsaturated | 47 |
| TG 47:4\|TG 13:1_16:1_18:2_[M+NH4]+_22.104 | LGR4/5dLKO HFD - LGR4/5dLKO ND | 3.370552796 | 1.04892E-05 | TG | Polyunsaturated | 47 |
| TG 52:4;1O\|TG 17:1_17:1_18:2;1O_[M+NH4]+_23.389 | LGR4/5dLKO HFD - LGR4/5dLKO ND | 3.401259164 | 0.00866123 | OxTG | Polyunsaturated | 52 |
| TG 52:4;1O\|TG 17:1_17:1_18:2;1O_[M+NH4]+_23.389 | LGR4/5dLKO HFD - LGR4/5dLKO ND | 3.401259164 | 0.00866123 | OxTG | Polyunsaturated | 52 |
| CE 24:3_[M+NH4]+_26.562 | LGR4/5dLKO HFD - LGR4/5dLKO ND | 3.470151906 | 0.007744885 | CE | Polyunsaturated | 24 |
| CE 24:3_[M+NH4]+_26.562 | LGR4/5dLKO HFD - LGR4/5dLKO ND | 3.470151906 | 0.007744885 | CE | Polyunsaturated | 24 |
| DG 40:6\|DG 18:1_22:5_[M+NH4]+_18.693 | LGR4/5dLKO HFD - LGR4/5dLKO ND | 3.496859652 | 3.3128E-09 | DG | Polyunsaturated | 40 |
| DG 40:6\|DG 18:1_22:5_[M+NH4]+_18.693 | LGR4/5dLKO HFD - LGR4/5dLKO ND | 3.496859652 | 3.3128E-09 | DG | Polyunsaturated | 40 |
| TG 54:5;1O\|TG 18:1_18:1_18:3;1O_[M+NH4]+_23.358 | LGR4/5dLKO HFD - LGR4/5dLKO ND | 3.527227257 | 0.000657152 | OxTG | Polyunsaturated | 54 |
| TG 54:5;1O\|TG 18:1_18:1_18:3;1O_[M+NH4]+_23.358 | LGR4/5dLKO HFD - LGR4/5dLKO ND | 3.527227257 | 0.000657152 | OxTG | Polyunsaturated | 54 |
| DG 37:2\|DG 18:1_19:1_[M+NH4]+_19.945 | LGR4/5dLKO HFD - LGR4/5dLKO ND | 3.715501322 | 2.39857E-08 | DG | Polyunsaturated | 37 |
| DG 37:2\|DG 18:1_19:1_[M+NH4]+_19.945 | LGR4/5dLKO HFD - LGR4/5dLKO ND | 3.715501322 | 2.39857E-08 | DG | Polyunsaturated | 37 |
| TG 54:5;1O\|TG 18:1_18:2_18:2;1O_[M+NH4]+_21.921 | LGR4/5dLKO HFD - LGR4/5dLKO ND | 3.87136679 | 0.008033922 | OxTG | Polyunsaturated | 54 |
| TG 54:5;1O\|TG 18:1_18:2_18:2;1O_[M+NH4]+_21.921 | LGR4/5dLKO HFD - LGR4/5dLKO ND | 3.87136679 | 0.008033922 | OxTG | Polyunsaturated | 54 |

| | | | | | | |
|---|---|---|---|---|---|---|
| TG 47:3\|TG 13:1_16:1_18:1_[M+NH4]+_22.949 | LGR4/5dLKO HFD - LGR4/5dLKO ND | 3.944109309 | 1.20418E-10 | TG | Polyunsaturated | 47 |
| TG 47:3\|TG 13:1_16:1_18:1_[M+NH4]+_22.949 | LGR4/5dLKO HFD - LGR4/5dLKO ND | 3.944109309 | 1.20418E-10 | TG | Polyunsaturated | 47 |
| TG 47:4\|TG 13:1_16:1_18:2_[M+NH4]+_22.305 | LGR4/5dLKO HFD - LGR4/5dLKO ND | 4.019356954 | 1.5133E-09 | TG | Polyunsaturated | 47 |
| TG 47:4\|TG 13:1_16:1_18:2_[M+NH4]+_22.305 | LGR4/5dLKO HFD - LGR4/5dLKO ND | 4.019356954 | 1.5133E-09 | TG | Polyunsaturated | 47 |
| TG 53:2\|TG 16:0_18:1_19:1_[M+NH4]+_25.752 | LGR4/5dLKO HFD - LGR4/5dLKO ND | 4.114938286 | 1.02792E-10 | TG | Polyunsaturated | 53 |
| TG 53:2\|TG 16:0_18:1_19:1_[M+NH4]+_25.752 | LGR4/5dLKO HFD - LGR4/5dLKO ND | 4.114938286 | 1.02792E-10 | TG | Polyunsaturated | 53 |
| TG 45:3\|TG 13:1_16:1_16:1_[M+NH4]+_22.288 | LGR4/5dLKO HFD - LGR4/5dLKO ND | 4.242826175 | 6.93322E-10 | TG | Polyunsaturated | 45 |
| TG 45:3\|TG 13:1_16:1_16:1_[M+NH4]+_22.288 | LGR4/5dLKO HFD - LGR4/5dLKO ND | 4.242826175 | 6.93322E-10 | TG | Polyunsaturated | 45 |
| DG 40:4\|DG 18:1_22:3_[M+NH4]+_19.89 | LGR4/5dLKO HFD - LGR4/5dLKO ND | 4.428406991 | 8.5696E-12 | DG | Polyunsaturated | 40 |
| DG 40:4\|DG 18:1_22:3_[M+NH4]+_19.89 | LGR4/5dLKO HFD - LGR4/5dLKO ND | 4.428406991 | 8.5696E-12 | DG | Polyunsaturated | 40 |
| TG 69:3;O2\|TG 16:0_18:1_18:1;O(FA 17:0)_[M+NH4]+_26.613 | LGR4/5dLKO HFD - LGR4/5dLKO ND | 4.562211414 | 0.000853475 | TG_EST | Polyunsaturated | 69 |
| TG 69:3;O2\|TG 16:0_18:1_18:1;O(FA 17:0)_[M+NH4]+_26.613 | LGR4/5dLKO HFD - LGR4/5dLKO ND | 4.562211414 | 0.000853475 | TG_EST | Polyunsaturated | 69 |
| TG 51:2\|TG 16:0_17:1_18:1_[M+NH4]+_25.26 | LGR4/5dLKO HFD - LGR4/5dLKO ND | 4.804937345 | 7.97189E-13 | TG | Polyunsaturated | 51 |
| TG 51:2\|TG 16:0_17:1_18:1_[M+NH4]+_25.26 | LGR4/5dLKO HFD - LGR4/5dLKO ND | 4.804937345 | 7.97189E-13 | TG | Polyunsaturated | 51 |
| TG 45:2\|TG 16:0_13:1_16:1_[M+NH4]+_22.953 | LGR4/5dLKO HFD - LGR4/5dLKO ND | 4.878012708 | 1.42487E-07 | TG | Polyunsaturated | 45 |
| TG 45:2\|TG 16:0_13:1_16:1_[M+NH4]+_22.953 | LGR4/5dLKO HFD - LGR4/5dLKO ND | 4.878012708 | 1.42487E-07 | TG | Polyunsaturated | 45 |
| CE 22:3_[M+NH4]+_26.11 | LGR4/5dLKO HFD - LGR4/5dLKO ND | 4.961671654 | 3.25345E-05 | CE | Polyunsaturated | 22 |
| CE 22:3_[M+NH4]+_26.11 | LGR4/5dLKO HFD - LGR4/5dLKO ND | 4.961671654 | 3.25345E-05 | CE | Polyunsaturated | 22 |
| DG 35:1\|DG 17:0_18:1_[M+NH4]+_19.727 | LGR4/5dLKO HFD - LGR4/5dLKO ND | 5.303349903 | 1.04512E-09 | DG | Monounsaturated | 35 |
| DG 35:1\|DG 17:0_18:1_[M+NH4]+_19.727 | LGR4/5dLKO HFD - LGR4/5dLKO ND | 5.303349903 | 1.04512E-09 | DG | Monounsaturated | 35 |
| TG 55:4;1O\|TG 18:1_18:2_19:1;1O_[M+NH4]+_22.66 | LGR4/5dLKO HFD - LGR4/5dLKO ND | 5.342345023 | 0.002702514 | OxTG | Polyunsaturated | 55 |
| TG 55:4;1O\|TG 18:1_18:2_19:1;1O_[M+NH4]+_22.66 | LGR4/5dLKO HFD - LGR4/5dLKO ND | 5.342345023 | 0.002702514 | OxTG | Polyunsaturated | 55 |
| TG 71:4;O2\|TG 18:1_18:1_18:1;O(FA 17:0)_[M+NH4]+_26.582 | LGR4/5dLKO HFD - LGR4/5dLKO ND | 5.38881613 | 0.001374385 | TG_EST | Polyunsaturated | 71 |
| TG 71:4;O2\|TG 18:1_18:1_18:1;O(FA 17:0)_[M+NH4]+_26.582 | LGR4/5dLKO HFD - LGR4/5dLKO ND | 5.38881613 | 0.001374385 | TG_EST | Polyunsaturated | 71 |

| | | | | | | |
|---|---|---|---|---|---|---|
| DG 39:6\|DG 17:0_22:6_[M+NH4]+_18.284 | LGR4/5dLKO HFD - LGR4/5dLKO ND | 6.829618064 | 6.46754E-09 | DG | Polyunsaturated | 39 |
| DG 39:6\|DG 17:0_22:6_[M+NH4]+_18.284 | LGR4/5dLKO HFD - LGR4/5dLKO ND | 6.829618064 | 6.46754E-09 | DG | Polyunsaturated | 39 |
| PG 38:1\|PG 20:0_18:1_[M-H]-_16.778 | LGR4/5dLKO HFD - LGR4/5dLKO ND | 10.83167721 | 3.66798E-06 | PG | Monounsaturated | 38 |
| PG 38:1\|PG 20:0_18:1_[M-H]-_16.778 | LGR4/5dLKO HFD - LGR4/5dLKO ND | 10.83167721 | 3.66798E-06 | PG | Monounsaturated | 38 |
